# Thermal acclimation fails to confer a carbon budget advantage to invasive species over natives

**DOI:** 10.1101/2025.07.21.665833

**Authors:** Thibaut Juillard, Christoph Bachofen, Marco Conedera, Mattéo Dumont, Jean-Marc Limousin, Arianna Milano, Gianni Boris Pezzatti, Alberto Vilagrosa, Charlotte Grossiord

## Abstract

Both native and invasive plants can adjust photosynthesis and respiration when exposed to warmer temperatures. However, it is uncertain if invasive plants are more plastic and exhibit higher acclimation to rising temperatures than native ones, a trait that could contribute to their invasive behavior in novel environments. We compared the capacity of a highly invasive palm in central Europe (*Trachycarpus fortunei*) and two native co-occurring species (*Ilex aquifolium* and *Tilia cordata*) to acclimate photosynthesis and respiration to air temperature changes using a two-year-long transplant experiment across Europe (mean temperatures ranging from 8.4 to 21.8°C). We measured the optimal temperature of photosynthesis (T_opt_), the assimilation at optimal temperature (A_opt_), the thermal breath of photosynthesis (T_80_), the respiration at 25°C (R_25_), the temperature sensitivity of respiration (Q_10_), and simulated the whole-plant carbon budget. For all species, T_opt_, A_opt,_ and T_80_ increased with warming, while R_25_ decreased in the native species and Q_10_ decreased in the invasive species only. Consequently, acclimation enhanced the carbon budget of the invasive and native plants in the warm and hot sites. The invasive palm had a similar or lower acclimation capacity than other species and a lower but constant carbon budget across the European temperature gradient. Our work reveals that not all invasive plants exhibit greater photosynthetic plasticity than native ones, suggesting that temperature-driven enhancement of their carbon budget may play a limited role in future invasion processes.

## 1. Introduction

Global warming promotes the spread of invasive plant species worldwide, representing one of the most significant threats to plant biodiversity (Walther *et al*., 2009; Liu *et al*., 2017; Pyšek *et al*., 2020). Among possible factors, long-lived invasive plants can outperform native species under a warmer climate as they tend to be adapted to broader air temperature (T_air_) ranges (*e*.*g*., Higgins and Richardson, 2014; Finch *et al*., 2021; Turbelin and Catford, 2021), can extend their growing seasons more significantly (*e*.*g*., Fridley, 2012; Juillard *et al*., 2024), and acclimate various functional traits more extensively (*e*.*g*. Davidson, Jennions and Nicotra, 2011; Gioria *et al*., 2023). Higher carbon (C) uptake and plant productivity following a better thermal acclimation of photosynthesis and respiration may, in turn, enhance species competitiveness (Duan *et al*., 2013; Yu *et al*., 2019; Grossiord *et al*., 2022). Still, few studies investigated photosynthetic and respiratory thermal acclimation in the context of species invasiveness (Verlinden and Nijs, 2010; Godoy, Valladares and Castro-Díez, 2011; Ripley *et al*., 2020), with none addressing if invasive plants are more plastic than native ones with increasing temperature. Yet, understanding and predicting these processes would help reduce uncertainties in climate-vegetation models, where acclimation is often simplified or fully neglected (Crous, Uddling and De Kauwe, 2022) and would allow finding appropriate conservation strategies for forests highly susceptible to invasion as a consequence of global warming.

Plants generally acclimate rapidly to higher T_air_ (within one month; *e*.*g*., Kattge and Knorr, 2007; Kumarathunge *et al*., 2019) by increasing their optimal temperature (T_opt_, *i*.*e*., the temperature at which the net assimilation (A_net_) reaches its maximum), its corresponding optimal net assimilation value (A_opt_), and their thermal breath (T_80_), which represents the temperature range where photosynthesis reaches >80 % of its maximum (Yamori, Hikosaka and Way, 2014), thereby maintaining C gain despite warmer air (Kruse *et al*., 2019; Vico *et al*., 2019; Choury *et al*., 2022). Thermal acclimation of photosynthesis also depends on stomatal conductance (Kruse *et al*., 2019; Kullberg, Slot and Feeley, 2023) and phenology (Grossiord *et al*., 2022), which vary interannually (Petrik *et al*., 2022; Didion-Gency *et al*., 2024), leading to significant uncertainties in plant’s thermal acclimation capacity. Shifts in A_opt_, T_opt_, and T_80_ are mainly driven by an increase in the maximum catalytic activity of Rubisco (V_Cmax_) and the maximum ratio of electron transport (J_max_). Further, V_Cmax_ and J_max_ are both limited by nitrogen availability, therefore, thermal acclimation of photosynthesis also depends on nitrogen allocation, for which invasive plants tend to be more plastic than native ones (Feng and Fu, 2008; Funk, Glenwinkel and Sack, 2013).

In parallel, acclimation to higher T_air_ can also involve lowering the rate of respiration at 25°C (R_25_) and the respiration yield every 10°C (Q_10_) (Atkin and Tjoelker, 2003; Atkin, Bruhn and Tjoelker, 2005), thereby limiting C loss at high temperatures as respiration increases exponentially (Atkin and Tjoelker, 2003; Way and Yamori, 2014; Crous, Uddling and De Kauwe, 2022). Plants with higher photosynthesis acclimation can show higher respiration acclimation (Dusenge, Duarte and Way, 2019), although respiration can also acclimate more extensively to T_air_ than assimilation (Campbell *et al*., 2007; Way and Oren, 2010; Crous, Uddling and De Kauwe, 2022). Just as for photosynthesis, the duration of exposure to changed temperatures can influence respiration acclimation. Typically, acclimation of R_25_ to a particular T_air_ occurs during tissue development, while Q_10_ varies more rapidly with seasonal changes in ambient T_air_ (Atkin, Bruhn and Tjoelker, 2005). Some studies found that respiration tends to acclimate universally among plant species (Crous, Uddling and De Kauwe, 2022), and others reported acclimation to be species-specific independently of biomes or functional groups (*e.g*., evergreen *vs*. deciduous) (Slot and Kitajima (2015)). However, respiratory rates are inversely proportional to the leaf nitrogen content and the specific leaf area (Lee, Reich and Bolstad, 2005; Xu, Schuster and Griffin, 2007), frequently found to be higher in invasive species than in native ones (Baruch and Goldstein, 1999; Leishman *et al*., 2007). Therefore, temperature-induced reduction of respiratory rates could favor invasive species at high temperatures, but this remains largely unknown.

In this study, we compared the acclimation of leaf photosynthesis and respiration of the invasive *Trachycarpus fortunei*, a palm native from Southeastern China, with two co-occurring natives growing in the southern Alps in a sub-mediterranean climate (*i*.*e*., the evergreen *Ilex aquifolium* and deciduous *Tilia cordata*). Since the 2000s, *T. fortunei* has been colonizing natural forests worldwide, impacting the regeneration of native species (Fehr *et al*., 2023). In the southern Alps, the spread of *T. fortunei* has been correlated with the rise in T_air_ since the 1970s (ΔMAT: +1.7°C) (Walther *et al*., 2007; Fehr *et al*., 2023). As such, *T. fortunei* is suspected to benefit from warmer temperatures to outcompete its native competitors, but whether this includes enhanced photosynthetic and respiratory thermal acclimation is unknown. Using a transplant experiment in five sites covering a large range of mean T_air_ across Europe, we measured the responses of photosynthesis and respiration to temperature over two years. A soil-plant-atmosphere continuum (SPAC) model was used to estimate the impact of temperature acclimation on the leaf and whole-plant C budget over an entire growing season. We hypothesized that (1) the invasive *T*. *fortunei* acclimates more extensively than the native species to increased T_air_, leading to higher A_opt_, T_opt_, and T_80_. Similarly, (2) the invasive *T*. *fortunei* displays lower R_25_ and Q_10_ than native species with higher T_air_ and that, as such, (3) *T. fortunei* assimilates more C than native species in warmer sites.

## 2. Materials and methods

### 2.1 Experimental design

We selected five sites across a temperature gradient in Europe for a transplant experiment: one reference site located in the sub-Mediterranean climate where *T. fortunei* is the most invasive in Europe (*i*.*e*., the lowlands of the southern slope of the Swiss Alps), and four sites with differences of respectively −6°, −3°, +3°, and +6°C mean summer T_air_ (from April to October) compared to the reference site in the last ten years (2010-2020). This allowed us to assess the acclimation of the photosynthetic and respiration properties along a large T_air_ gradient, thereby covering regions where *T. fortunei* already has or could become invasive in future years (Fehr *et al*., 2023). In addition to the reference site in southern Switzerland (sub-Mediterranean climate), sites spanned from south-eastern Spain (thermic semi-arid), southern France (Mediterranean) to the Swiss Plateau (temperate), and the Swiss Alps (cold) (Fig. 1). The elevation of the sites ranged between 1 and 965 m a.s.l. (see Table 1 for more details). During the measurement period (2022 and 2023), mean annual T_air_ were 8.4, 13.3, 15.2, 17.7, and 21.8°C from the coldest to the warmest site, respectively, with slightly warmer conditions in 2022 than in 2023 (Fig. S1).

**Fig. 1:**
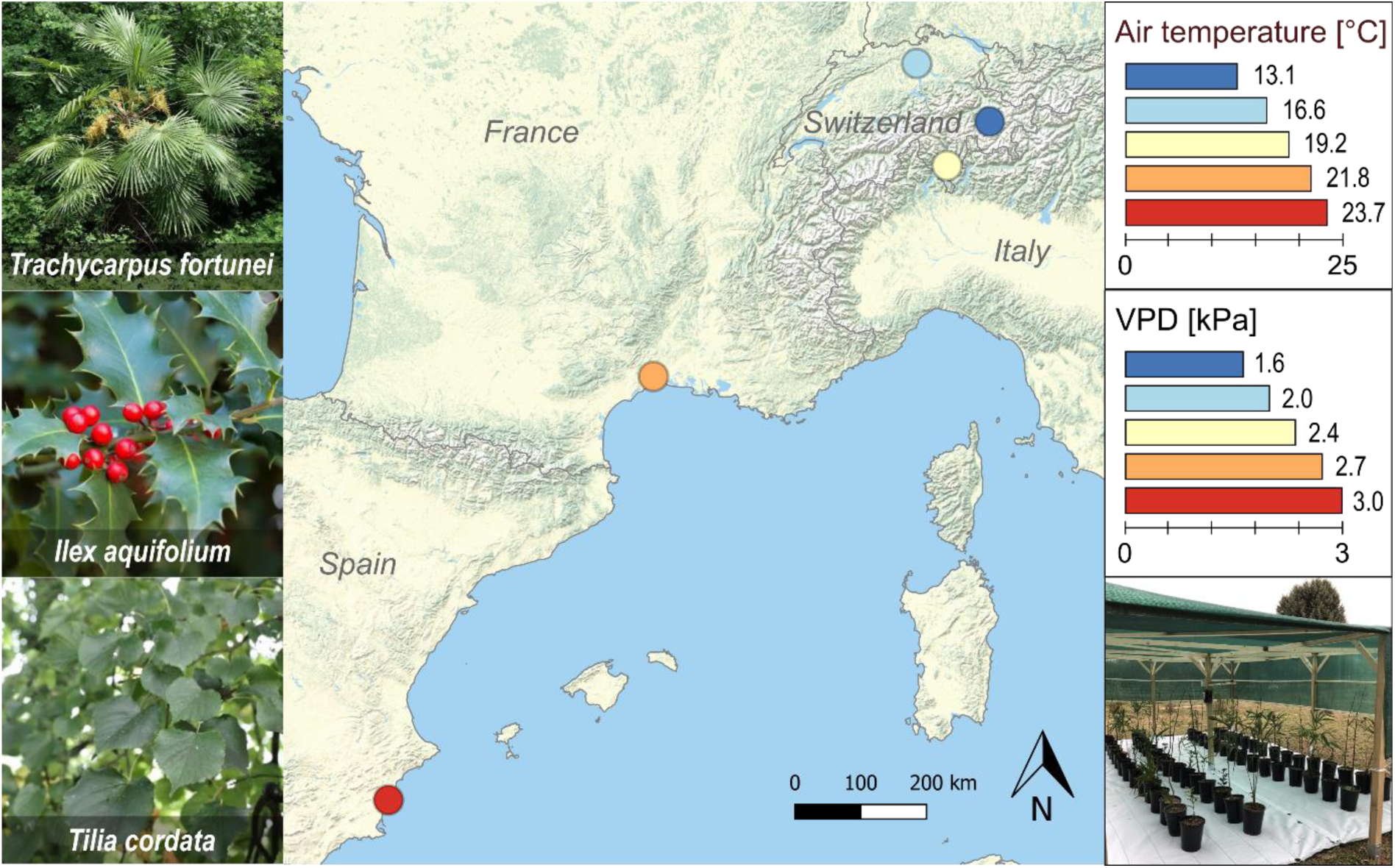
Map of the five experimental sites in Europe (from the coldest to the warmest: Filisur (cold, dark blue), Birmensdorf (temperate, light blue), Cadenazzo (sub-Mediterranean, yellow), Montpellier (Mediterranean, orange), and Guardamar del Segura (semi-arid, red). On the left are pictures of the three focal species included in the study. The right panels show the mean air temperature and VPD within each shading infrastructure from April to October 2022 and 2023. At the bottom right is a picture showing the shading infrastructure.

**Table 1:** Location of the experimental sites and associated climatic conditions.

| Experimental site | Filisur<br>(CH) | Birmensdorf<br>(CH) | Cadenazzo<br>(CH) | Montpellier<br>(FR) | Guardamar del<br>Segura (ES) |
| --- | --- | --- | --- | --- | --- |
| Climate | Cold | Temperate | Sub-Mediterranean | Mediterranean | Semi-arid |
| Location | 46°40'13.0"N | 47°21'41.6"N | 46°09'38.2"N | 43°38'19.2"N | 38°05'46.2"N |
|  | 9°40'06.6"E | 8°27'21.5"E | 8°56'00.9"E | 3°51'41.1"E | 0°38'51.7"W |
| Altitude [m a.s.l.] | 965 | 540 | 203 | 57 | 1 |
| MAT* 2010-2020 [°C] | 8.4 | 12.0 | 12.3 | 15.8 | 18.7 |
| T<sub>air</sub>* April-October 2010-2020 [°C] | 12.6 | 16.5 | 17.7 | 20.1 | 22.5 |

\*MAT = mean annual temperature; T<sub>air</sub> = air temperature

*T. fortunei* typically co-occurs with other species in the forest understory (*e*.*g*., the native evergreen *Ilex aquifolium* and deciduous *Tilia cordata*), which it occasionally excludes when dominating the canopy (Fehr *et al*., 2023). To simulate the natural half-shady understory environment, we constructed a wooden shading infrastructure (6 x 4 x 1.8 m) in an unobstructed area on all sites. The top and sides were covered with a permeable shading tissue obstructing 70% of sunlight. Mean yearly incoming solar radiation for 2022-2023 was similar among sites (Fig. S1).

In November 2021, we dug out 200 saplings of *T. fortunei* of 50-70 cm in height in a freshly invaded sub-Mediterranean forest (Cugnasco, Switzerland; 46°10’15’’ N, 8°55’49’’ E, 205 m a.s.l., MAT 12.2 °C, MAP: 1757 mm; 1.1 km from our reference site). We further bought 200 saplings of *T. cordata* (Morbio Superiore, Switzerland) and 150 saplings of *I. aquifolium* (Wiler bei Utzenstorf, Switzerland) from commercial producers. All individuals were immediately potted in 7 L pots with generic sandy forest soil made of 20% peat and mineral substrate (pH = 6.3; Ökohum; DE) and overwintered at the reference site.

In March 2022, 100 healthy individuals of each species (*i*.*e*., 300 saplings) were randomly separated into five groups of 20 individuals per species and transported to the five sites (n = 60 saplings per site). The plants for the coldest site were kept at the second coldest one (temperate climate) until April 2022 and from December 2022 to March 2023 to avoid exposure to temperatures below −15°C as it could have killed *T. fortunei* (Fehr and Burga, 2016). Despite this, 75% of the individuals of *T. fortunei* at the two coldest sites suffered frost damage during the winter of 2022-2023, limiting the number of replicates to four in 2023. At each site, pots were placed inside the shading infrastructure in four rows of 15 individuals each, alternating species to randomize potential differences in light and wind conditions. Each pot was individually connected to a drip irrigation system equipped with individual pressure resistances (Allenspach GreenTech AG, CH) to guarantee the same irrigation for each pot. Plants were watered automatically every two days at dawn to field capacity to ensure water availability throughout the experiment. We recorded T_air_, RH, and solar radiation (data logger: EM-100; air temperature and humidity: Atmos-14, Meter Group Inc.; USA; light quantum sensor: SQ-100X-SS, Apogee Instruments; USA) inside each shading infrastructure every 30 minutes at 1,5 m above the ground. We also monitored the soil moisture of each plant’s pot at 10 cm depth monthly with a soil moisture meter (TDR-100, Spectrum Technologies; USA).

For two years (2022-2023), the measurement campaigns were conducted in May, July, and September (six in total) during two consecutive growing seasons (*i*.*e*., 2022 and 2023). We tracked leaf flush and senescence at each site with phenocams (IPX5, VisorTech; DE). To guarantee that we measured fully expanded leaves, the first campaign of each year in May was conducted one month after all individuals of *T. cordata* had flushed.

### 2.2 CO_2_ assimilation and respiration responses to air temperature

We measured the optimal temperature (T_opt_), net assimilation at the optimal temperature (A_opt_), thermal breath (T_80_), dark respiration at 25°C (R_25_), and respiration yield (Q_10_) of one-year-old (or fully developed of the current year in deciduous *T. cordata*), undamaged leaves of 10 individuals per species (4 individuals of *T. fortunei* in 2023). Hence, every year in May, one leaf per individual was selected for T_opt_, A_opt_, and T_80_ measurements, while an adjacent leaf was chosen for R_25,_ and Q_10._

T_opt_, A_opt_, and T_80_ were obtained through CO_2_ assimilation response to T_air_ curves using portable photosynthesis systems (Li-6800, Licor Biosciences; USA) similar to Gauthey *et al*., (2023) and Deluigi *et al.,* (2025). Measurements were conducted approx. every 1-2 hours at five or six time points during the day, reflecting different ambient T_air_ from the sunrise (*i.e.*, the coldest T_air_ of the day) to the middle of the afternoon (*i.e.*, the warmest T_air_ of the day) to obtain the possible largest T_air_ range (from 12 to 35°C on average across sites and campaigns). Air temperature within the Li-6800 chamber was set according to the ambient T_air_, which was measured continuously (RS-91, RS Instruments; DE). By doing so, we ensured to track the temperature response curve of the ambient T_air_ and to avoid possible artifacts associated with differences between the ambient temperature experienced by the plants and the conditions in the cuvette, as well as bias associated with the calculations of T_L_ within gas exchange systems (Still *et al*., 2019). To extend the temperature range, we additionally decreased and increased the temperature of the cuvette by 5°C during the coldest and warmest time of the day, respectively. Relative humidity within the cuvette was increased at high temperatures to avoid stomatal closure due to high VPD. Hence, VPD ranged between 1 (fixed minimum) and maximum 3,5 kPa (when T_air_ reached > 38°C). T_opt_, A_opt_, and T_80_ were measured at saturating light (PPFD of 1500 μmol m^−2^ s^−1^) and ambient CO_2_ (400 ppm). Measurements of R_25_ and Q_10_ were conducted as described above, except that the leaf of each individual was wrapped in aluminum foil for at least 30 minutes before each measurement to dark-acclimate the leaves and that light was reduced to 0 μmol m^−2^ s^−1^ inside the cuvette during the measurements. All measurements were taken after the gas exchange rates had stabilized for at least 5 min.

We extracted T_opt_, A_opt_, and T_80_ of each individual by fitting our measurements of assimilation at different T_air_ with a parabolic curve in R (4.1.1, R Core Team, 2021), using the following equation as in Choury *et al*., (2022):

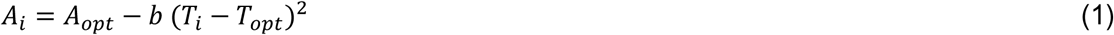

where A_i_ is the net assimilation at temperature T_i_ and b is the width of the curve. T_80_ was then calculated after resolving equation (1) for A_i_ = 0.8 A_opt_ and isolating T_i_. Similarly, R_25_ and Q_10_ were obtained after fitting the respiration measurements at different T_air_ with an exponential curve in R, using the following equation (Choury *et al*., 2022):

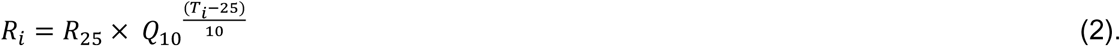

### 2.3 SPAC modeling of leaf-level photosynthesis and respiration

To assess the temperature effect on the annual leaf-level C uptake (C_leaf_), we modeled the instantaneous leaf-level net photosynthesis at 30-minute intervals from leaf flushing to senescence with a mechanistic soil-plant-atmosphere continuum (SPAC) model proposed by Garcia-Tejera *et al*. (2017). The model was modified to account for the acclimation of temperature responses of photosynthesis and respiration (Fig. 2). The original model calculates iteratively net assimilation (A_net_) and stomatal conductance (g_s_) based on environmental drivers and physiological parameters. A_net_ is calculated as:

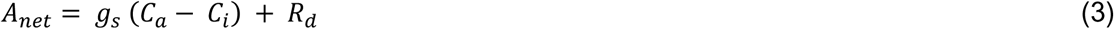

where C_a_ is the ambient CO_2_ concentration, C_i_ is the intracellular CO_2_ concentration, and R_d_ is the dark respiration rate. g_s_ is derived from its theoretical value at unlimited water availability (g_s,max_):

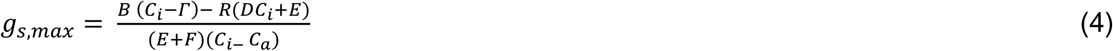

where B is the CO_2_ uptake limiting rate of either J_max_ or V_Cmax_, Γ is the CO_2_ compensation point of photosynthesis, E is a metric of carboxylation and oxygenation rates, and D and F are constants. g_s_ is then derived from g_s,max_ based on hydraulic parameters, water transport from the soil to the leaf, and transpiration (Fig. S2, equations in Tables S1 & S2).

**Fig. 2:**
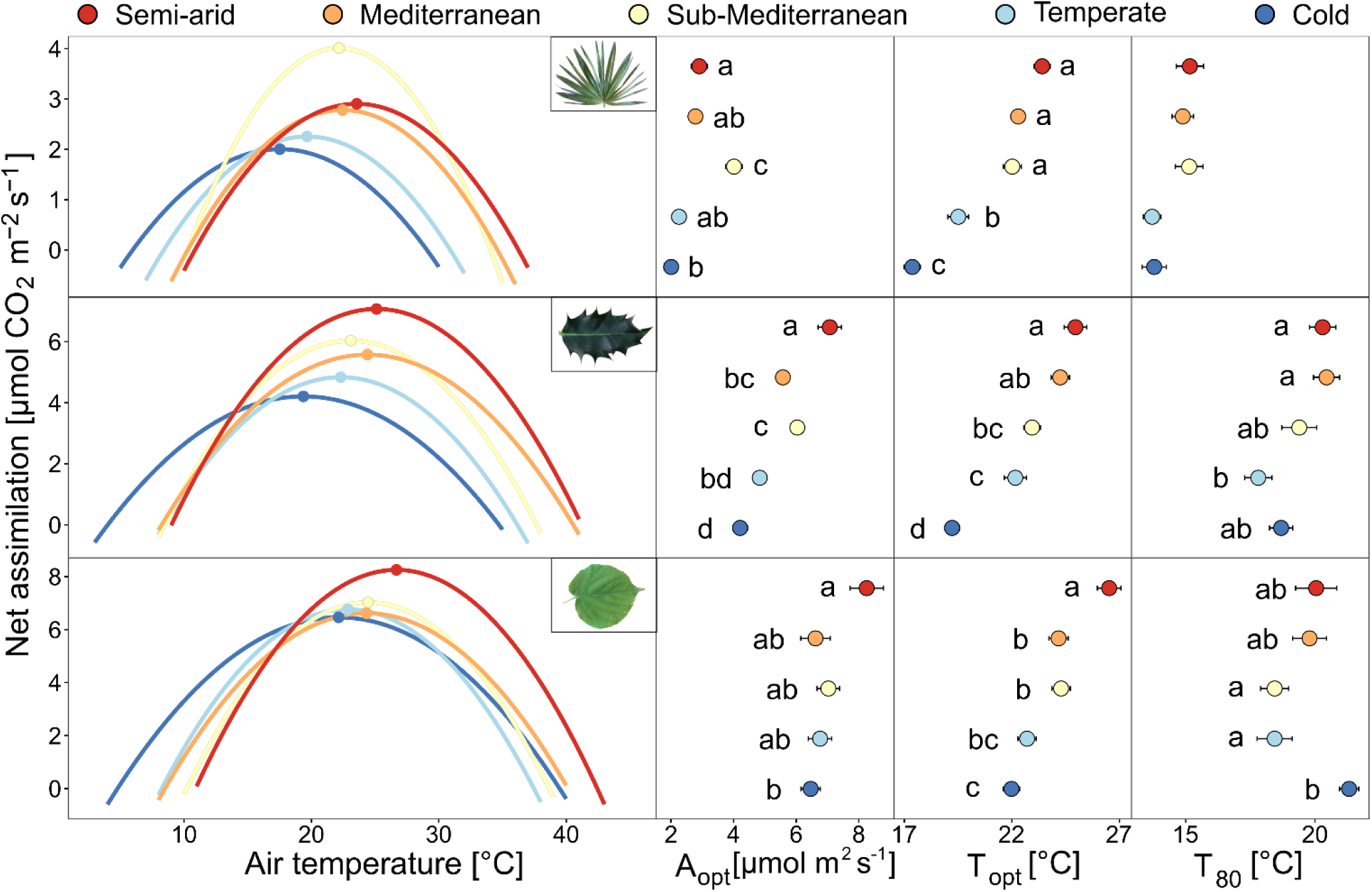
Assimilation-temperature response curves for the summer months (mean of May-October) of both years (2022-2023) for *T. fortunei*, *I. aquifolium*, and *T. cordata* at the five experimental sites (indicated with colors from blue to red going from the coldest to the warmest). On the right panels, the mean and standard errors of A_opt_, T_opt_, and T_80_ at each site are shown (n = 4-10 individuals per species). Different letters indicate significative differences (*p* < 0.05) between sites for each species.

To model leaf temperature (T_L_), we used a leaf energy balance model by Kevin Tu (http://landflux.org/Tools.php) as in Marias, Meinzer and Still (2017). As such, T_L_ varied with T_air_, light, RH, and g_s_ following the equations from Jones (1992), Sridhar and Elliott (2002), and Monteith and Reifsnyder (2007) (Table S1). As T_L_ impacts g_s_, and vice versa, we nested the SPAC optimization process for g_s_ calculations into a second optimization process to calculate T_L_ (Fig. 2). R_d_ was calculated with T_L_ following equation (2). V_Cmax_ and J_max_ were adjusted for T_L_ following the peaked Arrhenius function equation as in Kumarathunge *et al*. (2019). To incorporate T_L_ acclimation in our model, we adjusted the entropy factor (ΔS, J mol^-1^ K^-1^) and the activation energy (Ha, kJ mol^-1^) in the peaked Arrhenius function with the general coefficients proposed by Kumarathunge *et al*. (2019) (Tables S2 & S3). For respiration, we adjusted R_25_ with T_air_ with linear species-specific equations that we derived from our results (Tables S1):

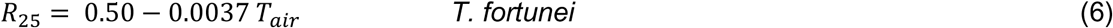

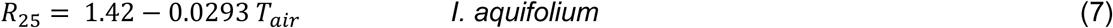

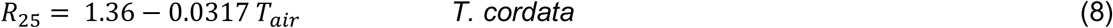

To parametrize the model, we measured J_max,25_ and V_c,max,25_ on five individuals of the three species in July 2023 at our reference site with the Li-6800 following the methodology used in Didion-Gency *et al*., (2022). In May 2022, we measured the minimum stomatal conductance (g_min_) on 4-5 saplings per species at EPFL (46°31’15.3"N, 6°34’04.0"E, Lausanne, CH). In September each year (2022 and 2023), we measured the leaf area of 4-10 individuals of each species at every site. For this, we photographed ten representative leaves of *I. aquifolium* and *T. cordata,* and all the leaves of *T. fortunei*. We extracted the leaf area using the software Fiji (Schindelin *et al*., 2012). The mean leaf area of *I. aquifolium* and *T. cordata* was multiplied by the total number of leaves of each individual to compute the total leaf area. Finally, the leaf area index (LAI) was obtained by dividing the total leaf area by the pot’s upper surface.

We calibrated and validated the model with A_net_ measurements that we took diurnally at the five experimental sites during every campaign. A_net_ was measured approximately every hour with the Li-6800 under ambient light, T_air_, and RH. We randomly selected 70 % of the measurements to calibrate influential model parameters that we could not measure similarly as in Grossiord *et al*. (2022) and Deluigi *et al*., (2025). The model was calibrated using a differential evolution (DEzs) Markov chain Monte Carlo (MCMC) sampler (Ter Braak and Vrugt, 2008) using the R package BayesianTools (Hartig *et al*., 2023). For each species, we run 10,000 iterations of three independent chains. We confirmed the convergence of the calibration with the Gelman–Rubin diagnostic and a threshold of 1.1 (Gelman and Rubin, 1992). We used the 30 % remaining measurements to assess the goodness-of-fit (RMSE, percent bias, and Nash-Sutcliff efficiency) between the measured and modeled A_net_ at each site. To estimate the effect of photosynthetic temperature acclimation on the plant C budget, we ran the model without acclimation by keeping V_C,max_, J_max_, R_25_, and Q_10_ constant (mean of the reference site).

### 2.4 Statistical analyses

Species and site differences in T_opt_, T_80_, A_opt_, R_25_, and Q_10_ were tested through analysis of variance by using species (*i*.*e*., *T. fortunei*, *I. aquifolium*, and *T. cordata*) and climate (semi-arid, Mediterranean, sub-Mediterranean, temperate, and cold) and their interaction as fixed effects. No variable transformation was required to ensure homoscedasticity. Tukey’s HSD post hoc tests were used to separately estimate differences between species or climates. Linear regressions were used to test the relationships between T_opt_, A_opt_, R_25_, Q_10,_ and T_air_ (mean of the two weeks preceding the measurements) and between A_opt_ and T_opt_. All statistical tests were done using the software R (4.1.1, R Core Team, 2021).

## 3. Results

### 3.1 Photosynthetic and respiration responses to air temperature

In both years, A_opt_, T_opt_, and T_80_ differed between species and climates (*p* < 0.001 for A_opt_ and T_opt_ and *p* < 0.05 for T_80_; Fig. 2 & Table S4), with generally higher values in the warmest site and lower values in the coldest one (Fig. S3 & Table S5). In all sites, A_opt_ and T_opt_ were lower in *T. fortunei* (2.86 µmol m^2^ s^-1^ and 21.3°C on average, respectively) than in *I. aquifolium* (5.5 µmol m^2^ s^-1^ and 22.7°C on average, respectively) and *T. cordata* (6.96 µmol m^2^ s^-1^ and 23.9°C on average, respectively), even in the reference sub-Mediterranean climate where *T. fortunei* is highly invasive (*p* < 0.001; Fig 2). Similarly, *T. fortunei* had a lower T_80_ (11.1°C on average) than *I. aquifolium* (13.9°C on average) and *T. cordata* (15.2°C on average) in all sites (Fig. 2).

A_opt_ and T_opt_ were strongly correlated with the mean T_air_ of the two preceding weeks, suggesting rapid photosynthetic acclimation (Fig. 3). The relationship between mean T_air_ and T_opt_ was significant for all species (*p* < 0.01), but steeper for the two evergreen species than for the deciduous one (+0.60, +0.55, and + 0.33°C T_opt_ per °C mean T_air_ for *T*. *fortunei*, *I*. *aquifolium*, and *T*. *cordata*, respectively). The same relationship with A_opt_ was significant in *T. fortunei* (+0.16 µmol m^2^ s^-1^ per °C; p < 0.001) and *I. aquifolium* (+0.19 µmol m^2^ s^-1^ per °C; *p* < 0.001), but not for *T. cordata* (Fig. 3). Air temperature and T_80_ were only correlated in *T. fortunei* (+ 0.19°C per °C air temperature, R^2^ = 0.25, *p* < 0.01; Fig. S4). A_opt_, T_opt_, and T_80_ were positively correlated in all species (*p* < 0.05; Fig. 4 & Fig. S5), except for T_opt_ and T_80_ in *T. fortunei*. In *T. fortunei*, A_opt_ increased by 0.12 µmol m^2^ s^-1^ per °C positive shift in T_opt_. The same correlation was steeper in *I. aquifolium* and *T. cordata* (+0.23 and +0.27 µmol m^2^ s^-1^ per °C shift in T_opt_, respectively; Fig. 4).

**Fig. 3:**
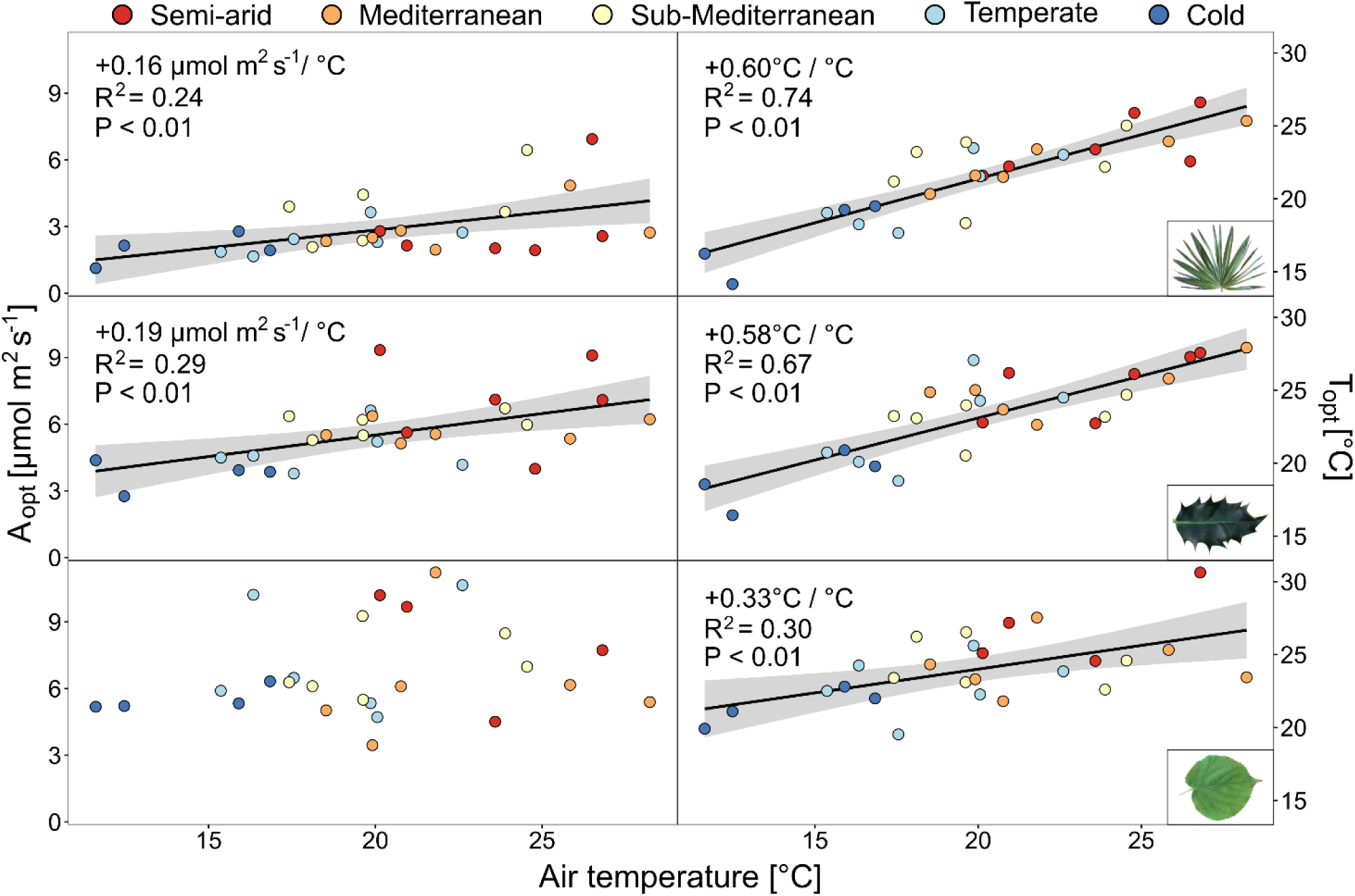
A_opt_ and T_opt_ (n = 4-10 individuals per species) averaged by campaigns in relation to air temperature of the two weeks preceding the measurements for *T*. *fortunei*, *I*. *aquifolium*, and *T*. *cordata*. Colors represent climates from blue to red, from the coldest to the warmest. Linear regression lines were added when significant.

**Fig. 4:**
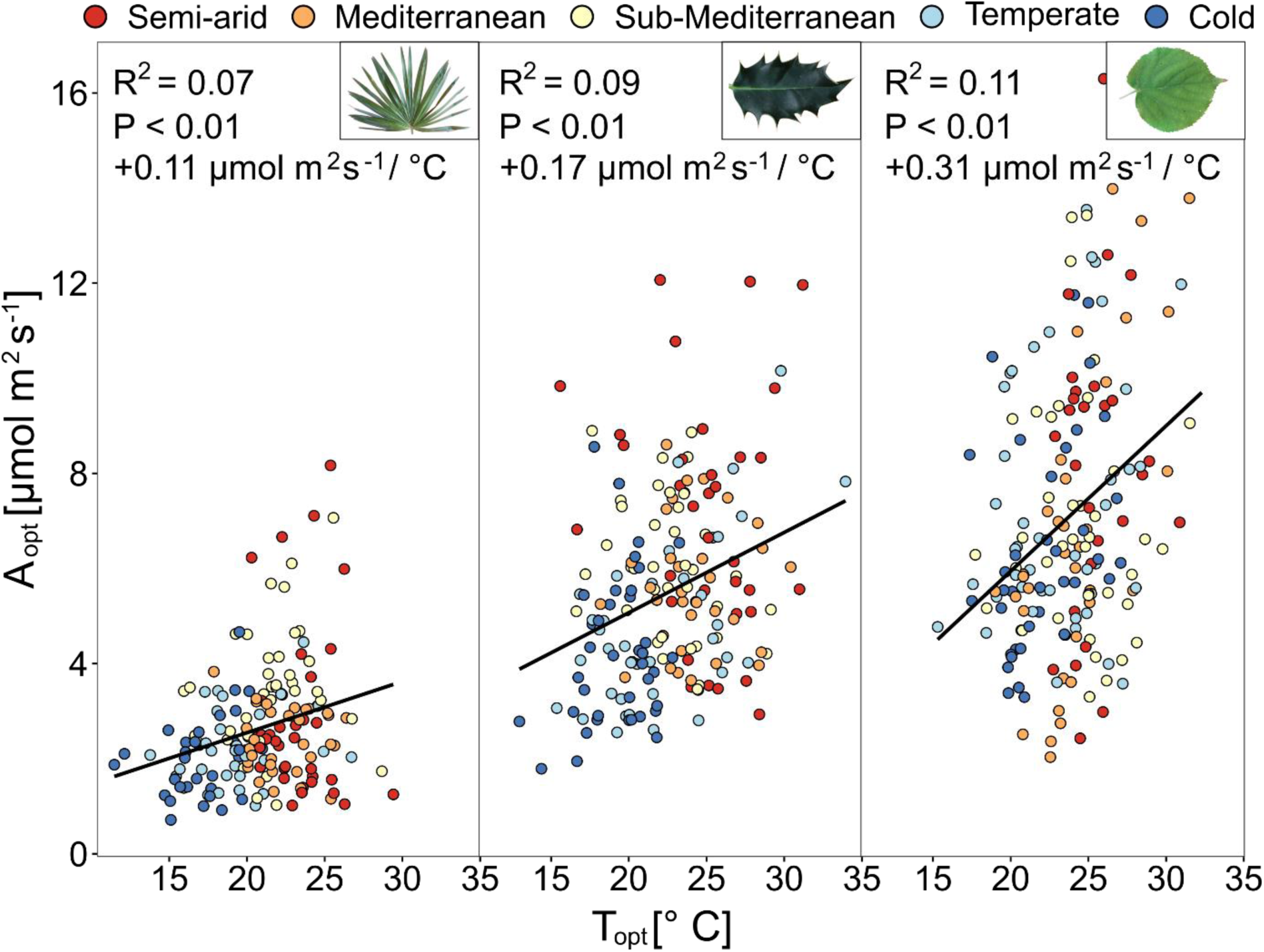
Assimilation at the optimal temperature (A_opt_) in function of the optimal temperature of photosynthesis (T_opt_) (n = 4-10 individuals per species) in *T. fortunei*, *I. aquifolium*, and *T. cordata* during all campaigns in 2022 and 2023. Colors represent sites from blue to red, from the coldest to the warmest. Linear regression lines were added when significant.

R_25_ varied across sites and species (*p* < 0.001; Fig. 5 & Table S4) with higher values in the colder sites, except in *T*. *fortunei*. On average, R_25_ was almost half as high in *T. fortunei* (0.40 µmol m^2^ s^-1^) than in *I. aquifolium* and *T. cordata* (0.85 and 0.75 µmol m^2^ s^-1^, respectively) (Fig. 5). R_25_ decreased with increasing mean T_air_ of the two preceding weeks in *I. aquifolium* and *T. cordata* (−0.03 µmol m^2^ s^-1^ / °C in both species, P < 0.01; Fig. 6) but not in *T. fortunei,* suggesting limited temperature acclimation in that species. On the other hand, while Q_10_ was different between species and sites (*p* < 0.001; Fig. 5 & Table S4), Q_10_ decreased with increasing mean T_air_ only in *T. fortunei* (Fig. 6).

**Fig. 5:**
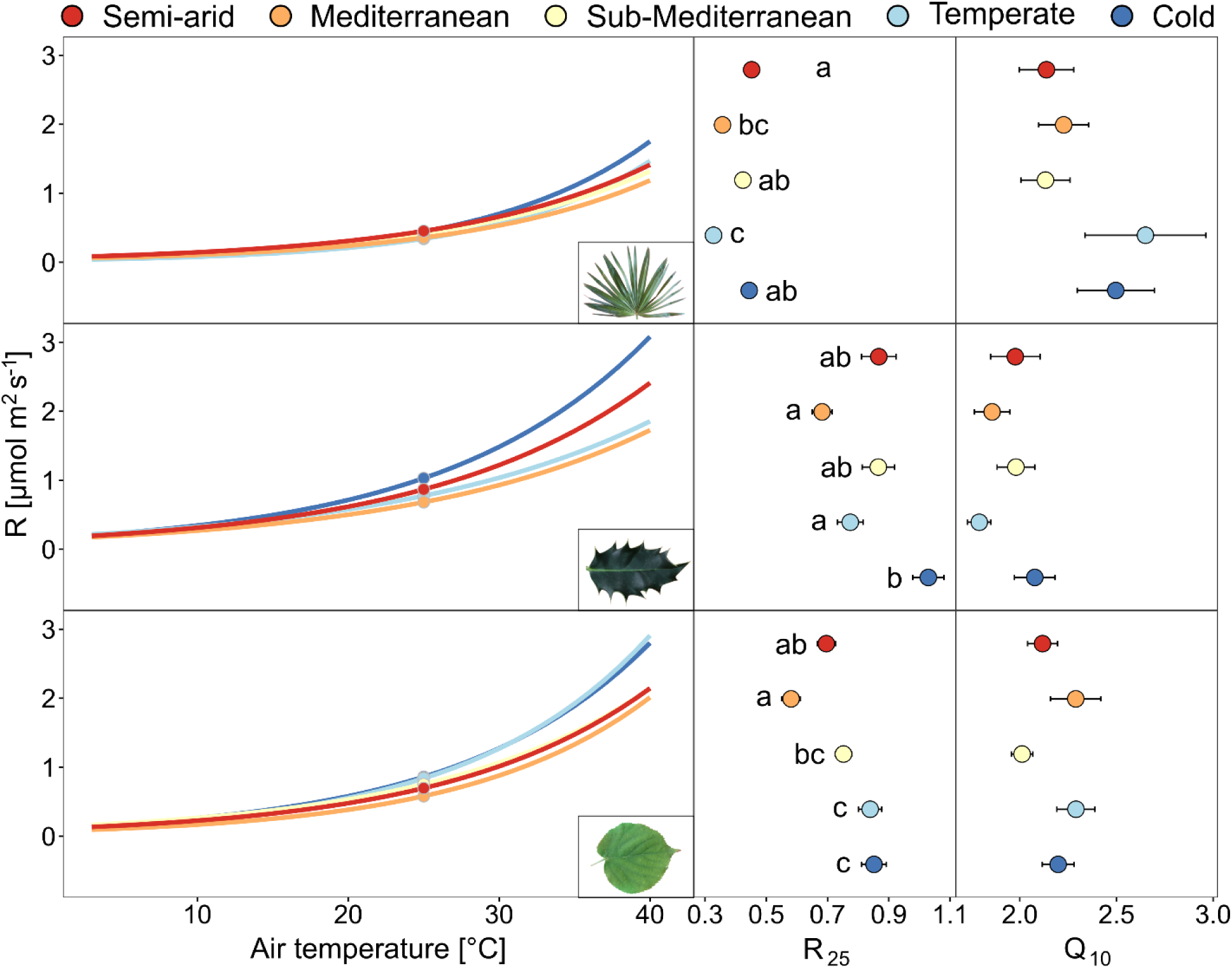
Respiration-temperature response curves for the summer months (mean of May-October) of both years (2022-2023) for *T. fortunei*, *I. aquifolium*, and *T. cordata* at the five experimental sites (indicated with colors from blue to red, from the coldest to the warmest). The right panels show the mean and standard errors of R_25_ and Q_10_ at each site (n = 4-10 individuals per species). Different letters indicate significant differences (*p* < 0.05) between the sites.

**Fig. 6:**
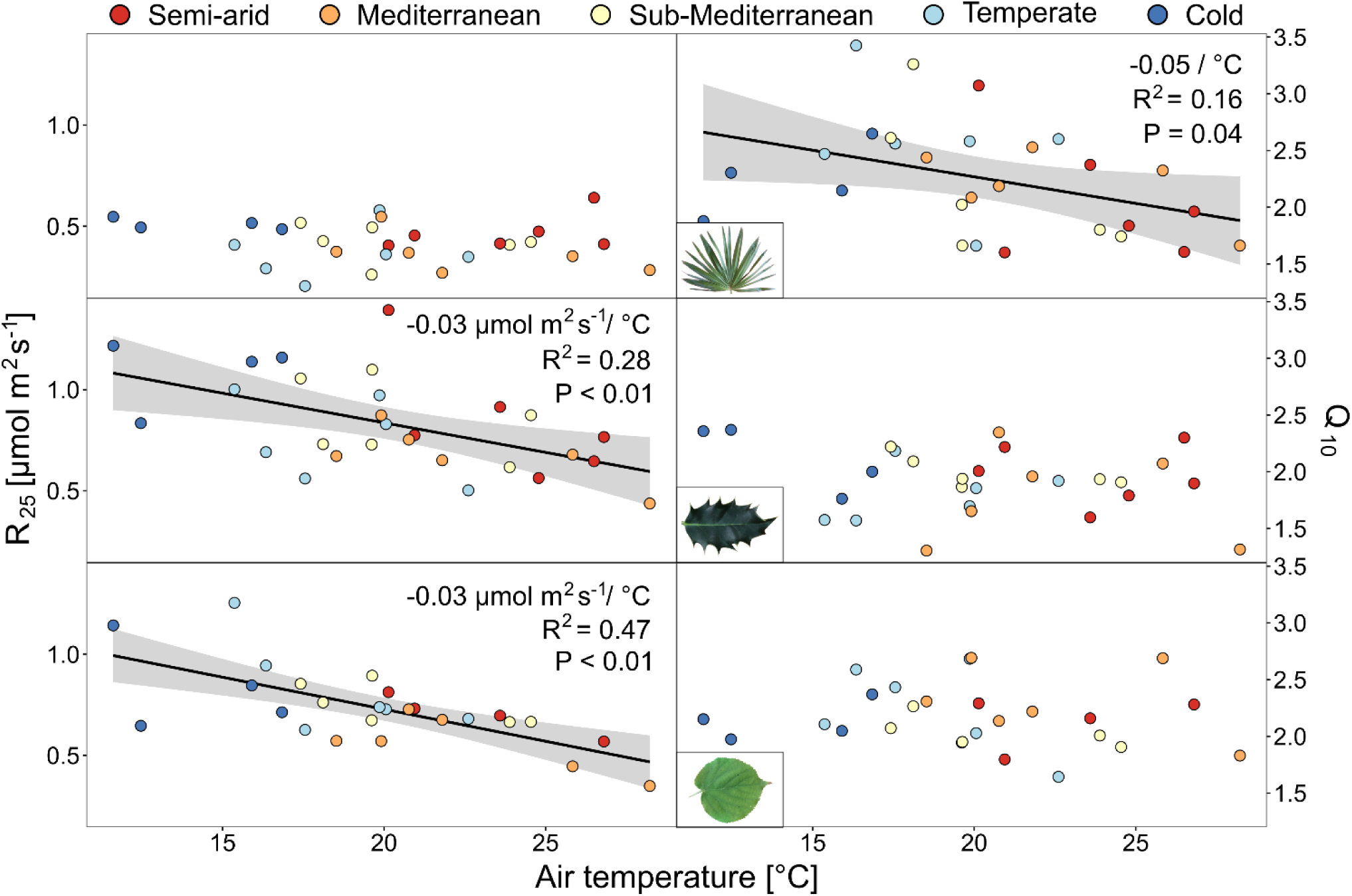
R_25_ and Q_10_ (n = 4-10 individuals per species) averaged by campaigns in relation to the air temperature of the two weeks preceding the measurements for *T. fortunei*, *I. aquifolium*, and *T. cordata*. Colors represent sites from blue to red, from the coldest to the warmest. Linear regression lines were added when significant.

### 3.2 Modelled carbon uptake

The SPAC model produced more precise predictions for *I. aquifolium* and *T. cordata* than for *T. fortunei* (R^2^ = 0.61, 0.66, and 0.17; NSE = 0.47, 0.62, and 0.11, respectively). Still, bias was low in all species (−0.61, 1.41, and 1.72%, respectively, Table S6), indicating a high accuracy of the model.

Overall, the model revealed significant differences in the annual leaf-level C uptake (C_leaf_) between species (*i*.*e*., 360, 345, and 587 gC m^-2^ on average for *T*. *fortunei*, *I*. *aquifolium*, and *T*. *cordata*, respectively; Fig. 7) and climates (*p* < 0.001), as well as an interaction between species and climate (*P* = 0.016, Table S7). At the reference site, *T. fortunei* had the lowest C_leaf_ (321 gC m^-2^) compared to *I. aquifolium* and *T*. *cordata* (418 and 615 gC m^-2^, respectively; Fig. 7). However, *T*. *fortunei* maintained a relatively constant C_leaf_ between the sites except at the warmest site, where a high R resulted in a lower C_leaf_ (Fig. 7 & Table S8). In contrast, both native species show the lowest C_leaf_ at the coldest site, a progressively higher C_leaf_ until a peak at the Mediterranean site, and a subsequent decrease at the warmest site (*i*.*e*., semi-arid) (Fig. 7). Notably, *T*. *cordata* was the only species that kept a similarly high C_leaf_ at the warmest site as at the reference sites despite higher R (Fig. 7).

**Fig. 7:**
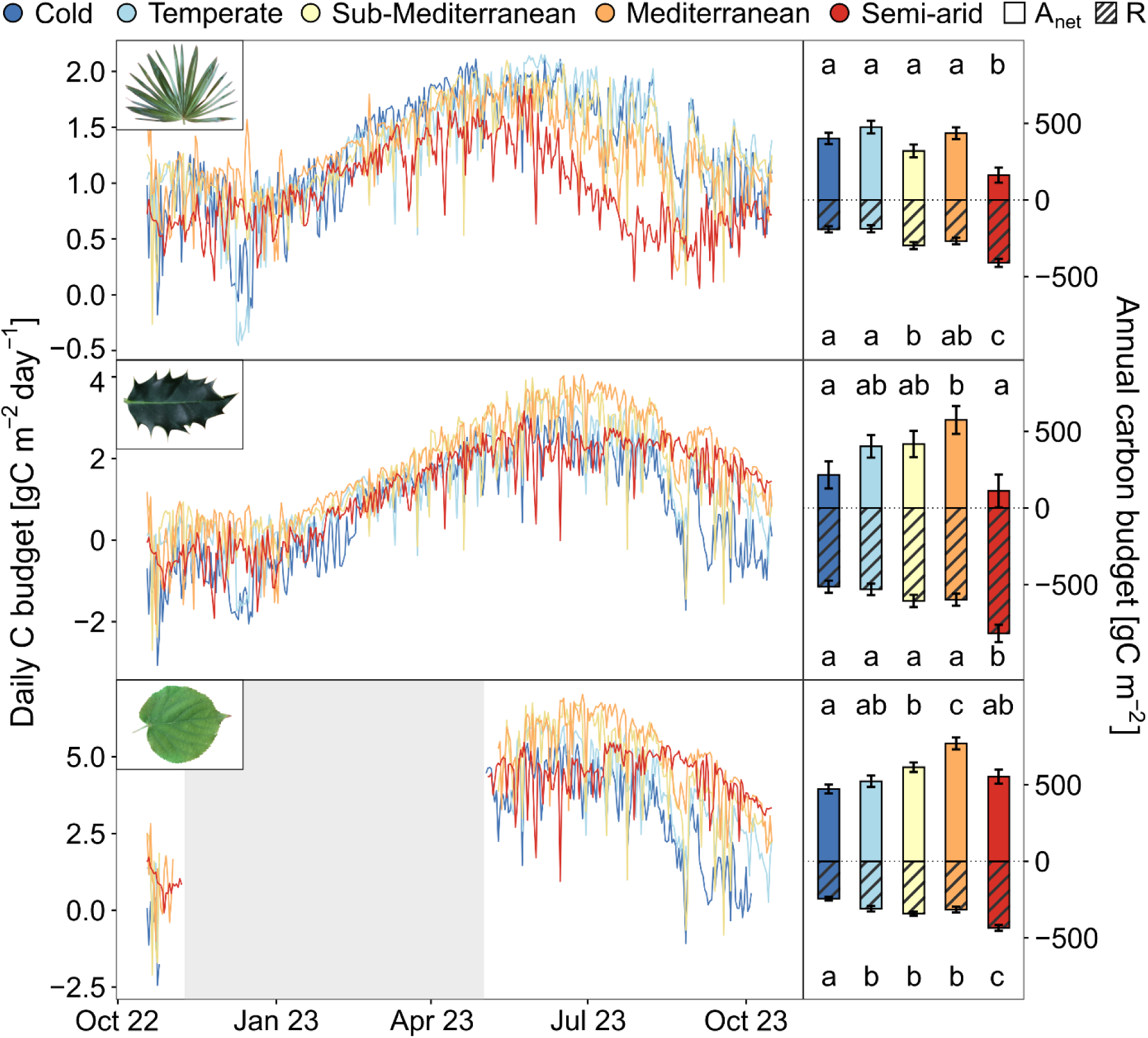
Daily mean leaf-level C budget in *T. fortunei*, *I. aquifolium*, and *T. cordata* at the five experimental sites from 15^th^ October 2022 to 15^th^ October 2023. The right panels show the annual leaf-level C uptake for each site over the same period. Net assimilation corresponds to the plain bar, whereas respiration is dashed. Error bars indicate the uncertainty of J_max,25_, V_Cmax,25_, R_25_, and Q_10_ (n = 37–57). Different letters indicate significant differences (*p* < 0.05) between the sites.

Total leaf area did not vary largely between sites, apart for *T*. *fortunei*, which had a larger leaf area in the three warmest sites compared to the two colder ones (*i*.*e*., because of frost damage in winter 2022-2023; Fig. S6). As a consequence, total C uptake (C_tot_) showed similar patterns as C_leaf_ across sites for all species (*i*.*e*., peaking at the Mediterranean site; Fig. S7) with higher C_tot_ in *T. cordata* (*i*.*e*., because of its higher leaf area; Fig. S6) compared to the other species. In contrast to C_leaf_, *T*. *fortunei* had lower C_tot_ in the two coldest sites because of reduced leaf area.

Acclimation of photosynthesis and respiration contributed strongly to the variation in C_leaf_. Without considering acclimation, C_leaf_ of *T*. *fortunei* would have decreased at the temperate and Mediterranean sites (*p* < 0.05; Fig. 8 and Table S9) and remained similar than the C_leaf_ with acclimation at more extreme climates (cold and semi-arid conditions). Similarly, *T*. *cordata* would have had a significantly lower C_leaf_ at the Mediterranean and semi-arid sites. Contrastingly, acclimation did not modify the C_leaf_ in *I. aquifolium* (Fig. 8).

**Fig. 8:**
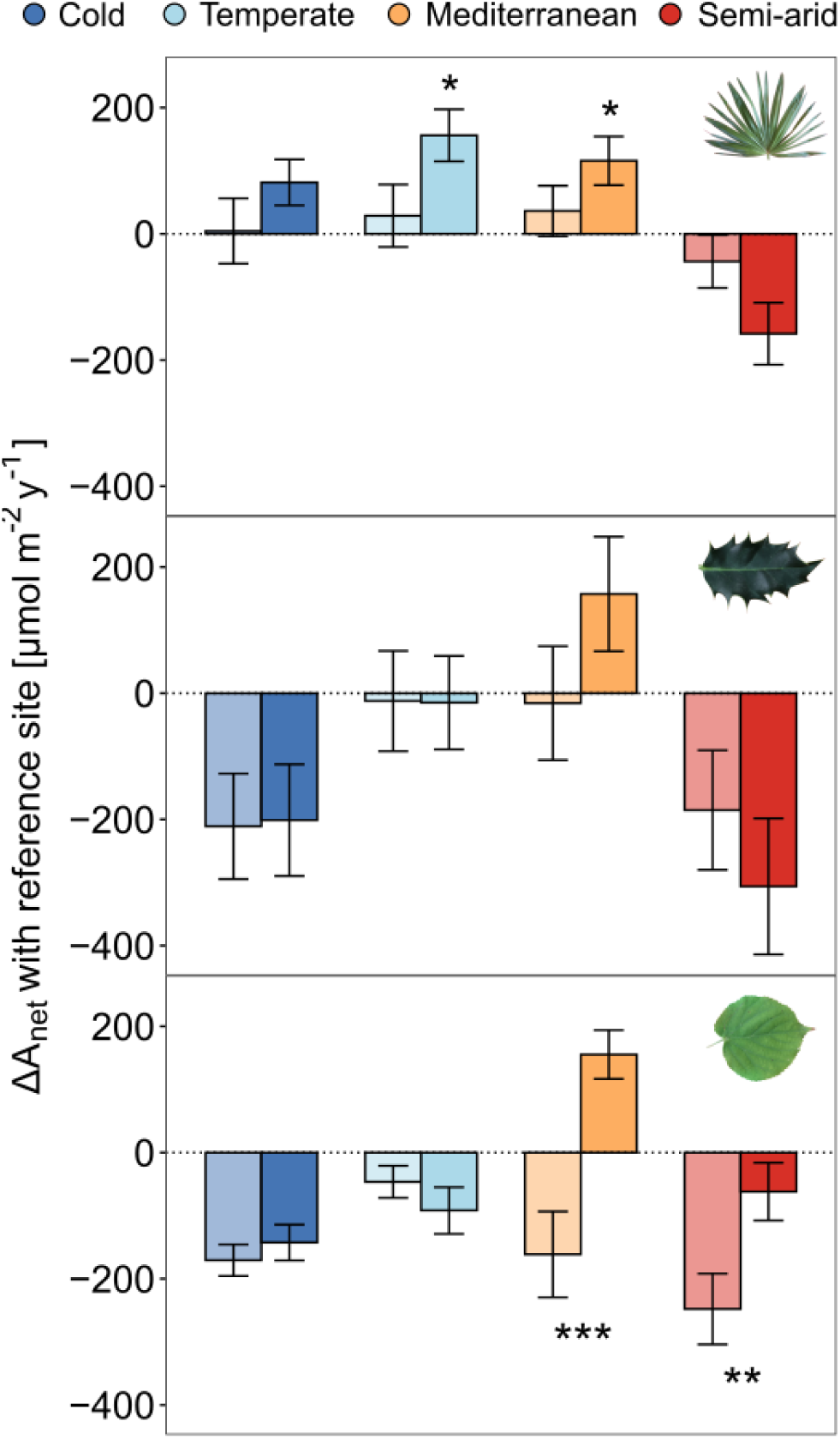
Differences between the modeled annual leaf-level C uptake (ΔA_net_) at every site and the reference site (sub-Mediterranean). Positive values indicate higher annual leaf-level C uptake in the respective sites compared to the reference one. For each site, the lighter bars show results obtained with non-acclimated physiological traits (*i*.*e*., the traits from the reference site), while the darker bars correspond to the outputs of the model with acclimated physiology. Error bars indicate the uncertainty of modeled J_max,25_ and V_Cmax,25_, R_25_, and Q_10_ (n = 37–57). Significant differences between acclimated and non-acclimated simulations are represented with stars (*P* < 0.05 = *; *P* < 0.01 = **; *P* < 0.001 = ***).

## 4. Discussion

### 4.1 Photosynthetic and respiratory acclimation is not higher in invasive palms than in native species

Against our initial hypothesis, the invasive *T*. *fortunei* did not show a higher photosynthetic acclimation than the native species, especially compared to the evergreen *I*. *aquifolium* that showed a similar thermal plasticity (Figs 3 & 4). All species had higher T_opt_ in the warmer sites than the colder ones (Fig. 3), but the two evergreen species showed similar T_opt_ acclimation rates at almost double that of the deciduous one (about +0.6°C *vs*. +0.3°C per degree T_air_; Fig 4.). Such differences between functional groups can be explained by the higher need for species with long-lasting tissues to tolerate a broader range of thermal conditions (Yamori, Hikosaka and Way, 2014). Despite those differences between functional groups, all our shifts in T_opt_ were within the range of +0.3 to 0.8 °C per degree T_air_, as found in most studies on leaf thermal acclimation (*e*.*g*., Kumarathunge *et al*., 2019; Choury *et al*., 2022; Crous, Uddling and De Kauwe, 2022). The T_opt_ shifts were probably driven by changes in the ratio between V_C,max_ and J_max_, as commonly found in other C3 species (Kattge and Knorr, 2007; Smith and Dukes, 2017), even if more work would be needed to confirm this.

In contrast to previous studies (Gunderson *et al*., 2010; Sendall, Lusk and Reich, 2016; Kruse *et al*., 2019), we also observed an increase of A_opt_ with rising T_air_ (Fig. 4) and generally higher values at warmer sites (Fig. 3). Several studies have underpinned the important role of stomatal aperture on A_opt_ shifts with temperature, especially compared to T_opt,_ which is generally less impacted by stomatal closure (Kruse *et al*., 2019; Akaji, Torimaru and Akada, 2024; Slot *et al*., 2024). In our study, plants were well-watered, and VPD was as low as technically possible during the measurements, allowing us to measure the thermal acclimation of A_opt_ mostly without the bias of increasing stomatal closure. These results are far-reaching as they suggest that VPD effects on stomata may hide the actual temperature acclimation potential of A_opt_ driven by higher enzymatic activity in warmer environments (Berry and Bjorkman, 1980). Still, similarly to T_opt_, *T*. *fortunei* did not show larger A_opt_ shifts than the other species (Fig. 3). Higher T_opt_ also led to higher A_opt_, and for every °C change in T_opt_, A_opt_ only increased by 0.11 µmol m^2^ s^-1^ in the invasive species, while it increased up to three times more for *T. cordata* (Fig. 4). However, *T*. *fortunei* was the only species increasing its T_80_ with T_air_ (+ 0.19°C per °C air temperature; Fig. S3). Slot and Winter (2017) found an increase in T_80_ that was accompanied by the rise in the J_max_:V_C,max_ ratio. Hence, a higher T_80_ could be either related to a higher J_max_, or a lower V_C,max_. The higher T_80_ in the invasive palm could thus indicate a decrease in V_C,max_ or increase in J_max_ with higher T_air_.

Contrary to our initial hypothesis, R_25_ in *T*. *fortunei* did not decrease with higher T_air_, showing no evidence of acclimation (Figs 5 & 6). Wang *et al*. (2020) observed that V_C,max_ and R_25_ were decreasing with increasing T_air_ as less active Rubisco is needed for a given value of V_c,max_ at higher temperatures, decreasing the respiratory costs for Rubisco turnover and therefore R_25_. As R_25_ of *T*. *fortunei* remained relatively constant across temperatures, it supports the notion that V_C,max_ remained stable as well in that species. The constant R_25_ may also indicate a lack of acclimation pressure, as at temperatures below 30°C, respiratory rates were low in this species (Fig. 5). This corroborates that palms can display lower foliar respiration rates than dicot species below 35 °C (Cavaleri, Oberbauer and Ryan, 2008). In contrast to *T*. *fortunei*, in the two native species, R_25_ decreased with higher T_air_ (−0.03 μmol CO_2_ m^−2^ s^−1^ per degree T_air_ on average; Fig. 6), with similar rates as in other studies (*e*.*g*., Reich *et al*., 2016; Zhu *et al*., 2020) following the consistent shifts of R_25_ with T_air_ found in a wide range of species and diverse experimental designs (Slot and Kitajima, 2015; Crous, Uddling and De Kauwe, 2022). Contrastingly, *T*. *fortunei* shifted Q_10_ while the native species did not (Figs 5 & 6). Similar acclimation of Q_10_ was found in other works focusing on evergreen species (*e*.*g*., Quan, Wang and Wang, 2020; Crous, Uddling and De Kauwe, 2022) and typically occurs when respiration is limited by low substrate availability (Atkin and Tjoelker, 2003; van de Weg, Fetcher and Shaver, 2013). The decrease in Q_10_ at high temperatures allowed the invasive palm to increase its A_net_ at warmer sites and could partially explain the broader T_80_ we observed at high temperatures (Fig. S3). Q_10_ shifts have been associated with short-term changes of respiration with T_air_ (Atkin and Tjoelker, 2003) and our findings contrast with studies measuring evergreen species in climatic chambers that observed no acclimation with temperature (*e*.*g*., Dusenge, Madhavji and Way, 2020; Choury *et al*., 2022), potentially because Q_10_ also varies seasonally and its acclimation depends on the warming duration and intensity (Atkin, Bruhn and Tjoelker, 2005; O’Sullivan *et al*., 2013). In fact, all our measured parameters (A_opt_, T_opt_, T_80_, R_25_, and Q_10_) shifted systematically with T_air_ within two weeks in the three species. These results align with other studies (*e.g*., Kattge and Knorr, 2007; Kumarathunge *et al*., 2019) stating that the commonly used 30-day acclimation period may be generally overestimated and that future works should consider using shorter acclimation periods (Smith and Dukes, 2013; Reich *et al*., 2016; Vico *et al*., 2019). Second, all species (*i*.*e*., invasive and native ones) demonstrated a short acclimation duration and the invasive palm did not benefit from a more extensive photosynthetic and respiratory acclimation than native tree species.

### 4.2 Impact of photosynthetic and respiratory acclimation on the leaf-level and plant C budget

Contrary to our expectations, the invasive palm generally showed lower C_leaf_ and C_tot_ than native species across all sites, even at the reference site where the palm invades the natural ecosystem (Figs. 7 and S6). While higher C gains increase the competitivity of plant species (Pearcy *et al*., 1987), lower C costs in invasive plants also play a significant role in invasion processes, and *T*. *fortunei* may have benefitted from other characteristics (*e*.*g*., longer photosynthetic period in autumn, lower whole-plant respiration, herbivory, and tissue turnover (Fridley *et al*., 2022; Juillard *et al*., 2024)) to be invasive under current climatic conditions. At the reference site, *T*. *fortunei* also shares these characteristics with several other natives (*e*.*g*., *Ilex aquifolium*, *Hedera helix*) and non-native evergreen species (*e*.*g*., *Prunus laurocerasus*) currently spreading in the understory of sub-Mediterranean deciduous forests, which could confirm the important role of these characteristics in the forest composition changes (Conedera *et al*., 2018).

The similar C_leaf_ of *T*. *fortunei* between the sites (Fig. 7) can be explained by the relatively low increase in A_opt_ per °C T_opt_ (+0.11 µmol m^2^ s^-1^ per °C T_opt_; Fig. 4), while T_opt_ showed the highest increase with T_air_ among the compared species (+0.60°C per °C T_air_). Hence, accurate A_opt_ measurements not biased by stomatal closure at high VPD are crucial to predict whole tree carbon exchange acclimation realistically. At the leaf level, the invasive palm performed similarly well at the reference site than in the two colder sites, while the native species had a lower C_leaf_ (Fig. 7), which shows that if *T*. *fortunei* is not damaged by frost in colder climates (Figs S6 & S7), its photosynthetic capacity allows to perform equally well as the native species. In contrast, native species acclimated more to the higher T_air_ at the Mediterranean site than the invasive palm (Fig 7 & S1), potentially limiting the capacity of the invasive palm to invade those environments. At the warmest site, C_leaf_ was lower in all species. Still, high temperatures especially limited the performance of *I*. *aquifolium* due to high respiratory rates (Figs 7 & 8), highlighting the critical role of respiration on the C budget at temperatures above 30°C.

Acclimation of photosynthesis and respiration allowed the invasive palm to increase C uptake and reduce C losses (leading to a higher C budget) in intermediate sites with milder conditions (Fig. 8). Nevertheless, the importance of acclimation was limited in extreme climates (both cold and hot ones), suggesting that shifts in photosynthetic and respiratory patterns are insufficient to compensate for the harsher climatic conditions. In contrast, acclimation allowed the native *T*. *cordata* to increase the C budget in the hottest environments (Fig. 8). Still, the highest C budget was observed in the Mediterranean climate (+2.5°C), suggesting a limited enhancement of the C budget under global warming. Previous work also reported increased C gains in some temperate trees with a rise between +2 and +5 °C T_air_, especially because warming also leads to a certain extent to a longer growing season in deciduous trees (Duan *et al*., 2013; Grossiord *et al*., 2022). On the other hand, the absence of photosynthetic and respiratory acclimation benefits for *I. aquifolium* (Fig. 8) could be explained by the high respiratory rates that drastically reduced the C_leaf_ in all sites, even the cold ones (Fig 7). As a shade-tolerant species, *I*. *aquifolium* tends to have high leaf N content, which has been associated with high respiration (Reich *et al*., 1998; Niinemets, Valladares and Ceulemans, 2003), potentially constraining the positive effect of T_air_ on its C budget.

Overall, our results suggest that photosynthetic and respiration thermal acclimation may not be the primary underlying mechanisms driving the fast propagation of the invasive palm in recent decades. Those findings are consistent with reports of invasive species conserving their initial niche (Verlinden and Nijs, 2010; Liu *et al*., 2020; Ripley *et al*., 2020). In addition, our results confirm that some long-lived invasive species have not reached their potential range yet, as observed in Bradley, Early and Sorte (2015). Their range expansion could, therefore, be predicted from physiological measurements directly but not from the extrapolation of their native or current range (Walther *et al*., 2009). Invasive plants often depend on many traits to colonize a new environment, and future invasion patterns in a warmer environment are difficult to predict. Our results suggest that low foliar respiratory rates may provide an advantage to the invasive palm over the native plants. Given that respiration plays a vital role at high temperatures, high respiratory rates often observed in fast-growing invasive species (Montesinos, 2022), could, therefore, reduce the C budget of fast-growing invasive species under higher temperatures.

## 5. Conclusion

Using a natural temperature gradient across European study sites, we showed that acclimation of photosynthesis and respiration rapidly occurs in young plants exposed to understory conditions and can play a significant role in the C budget of both invasive and native plant species. Contrary to our expectations, the invasive palm *T. fortunei* did not demonstrate higher acclimation capacity than its native competitors and had a lower (but more constant) C budget overall. In central and southern Europe, T_air_ during summer could increase by 5–7°C by the end of the century compared to 1995–2014 (SSP5-8.5) (Carvalho, Cardoso Pereira and Rocha, 2021), which would decrease the C budget of all species measured in this study. It is, therefore, likely that native sub-Mediterranean species will be outcompeted by more thermophilic invasive species in the future (Walther *et al*., 2009; Liu *et al*., 2017), which will likely lead to species composition changes in these forests.

## Data availability statement

Data will be made available on the Dryad Digital Repository.

## Acknowledgments

This study was supported by the Sandoz Family Foundation. CG and CB were supported by the Swiss National Science Foundation (310030_204697). AV was supported by EVER project (CIPROM/2022/37-Prometeu program, GVA). CEAM is funded by Generalitat Valenciana. We thank Janisse Deluigi, Stéphane Jenni, Samuel Reyes, Alex Tunas Corzon, Gabor Reiss, Luis Pizarro Pietro, Eugénie Mas, Kevin Knecht, Elias Ackermann, Omar Basquet, Alice Gauthey, Krzysztof Wroński, Jolan Wicht, Margaux Didion-Gency, Valentin Meister, Patrick Favre, and Maxwell Bergström for their precious help during the management of the experimental sites and the fieldwork. We also thank Mercedes Carmona and Diana Fernández in the Forestry nursery of Guardamar del Segura (Conselleria de Medio Ambiente, Infraestructuras y Territorio, Generalitat Valenciana, ES;), *Schutz Filisur Alpin Gartencenter* at Filisur (CH), and the CEFE CNRS at Montpellier (FR) for hosting the experimental sites and providing water access all along the experiment.

## Author Contributions

TJ, CG, and CB planned and designed the research. All authors carried out the fieldwork. TJ, CB, and MD performed data analyses. TJ wrote the manuscript draft. All authors contributed to the review and editing of the manuscript.

## References

Akaji, Y., Torimaru, T. and Akada, S. (2024) ‘Intraspecific variation in photosynthetic thermal acclimation in Fagus crenata seedlings from two populations growing at different elevations in northern Japan’, Tree Physiology, 44(8). doi: 10.1093/treephys/tpae093.

Atkin, O. K., Bruhn, D. and Tjoelker, M. G. (2005) Response of Plant Respiration to Changes in Temperature: Mechanisms and Consequences of Variations in Q10 Values and Acclimation, Advances in Photosynthesis and Respiration. doi: 10.1146/annurev.ph.37.030175.001511.

Atkin, O. K. and Tjoelker, M. G. (2003) ‘Thermal acclimation and the dynamic response of plant respiration to temperature’, Trends in Plant Science, 8(7), pp. 343–351. doi: 10.1016/S1360-1385(03)00136-5.

Baruch, Z. and Goldstein, G. (1999) ‘Leaf construction cost, nutrient concentration, and net CO2 assimilation of native and invasive species in Hawaii’, Oecologia, 121(2), pp. 183–192. doi: 10.1007/s004420050920.

Berry, J. and Bjorkman, O. (1980) ‘Photosynthetic Response and Adaptation to Temperature in Higher Plants’, Annual Review of Plant Physiology, 31(1), pp. 491–543. doi: 10.1146/annurev.pp.31.060180.002423.

Ter Braak, C. J. F. and Vrugt, J. A. (2008) ‘Differential Evolution Markov Chain with snooker updater and fewer chains’, Statistics and Computing, 18(4), pp. 435–446. doi: 10.1007/s11222-008-9104-9.

Bradley, B. A., Early, R. and Sorte, C. J. B. (2015) ‘Space to invade? Comparative range infilling and potential range of invasive and native plants’, Global Ecology and Biogeography, 24(3), pp. 348–359. doi: 10.1111/geb.12275.

Campbell, C. et al. (2007) ‘Acclimation of photosynthesis and respiration is asynchronous in response to changes in temperature regardless of plant functional group’, New Phytologist, 176(2), pp. 375–389. doi: 10.1111/j.1469-8137.2007.02183.x.

Carvalho, D., Cardoso Pereira, S. and Rocha, A. (2021) ‘Future surface temperatures over Europe according to CMIP6 climate projections: an analysis with original and bias-corrected data’, Climatic Change, 167(1–2), pp. 1–17. doi: 10.1007/s10584-021-03159-0.

Cavaleri, M. A., Oberbauer, S. F. and Ryan, M. G. (2008) ‘Foliar and ecosystem respiration in an old-growth tropical rain forest’, Plant, Cell and Environment, 31(4), pp. 473–483. doi: 10.1111/j.1365-3040.2008.01775.x.

Choury, Z. et al. (2022) ‘Tropical rainforest species have larger increases in temperature optima with warming than warm-temperate rainforest trees’, New Phytologist, 234(4), pp. 1220–1236. doi: 10.1111/nph.18077.

Conedera, M. et al. (2018) ‘Drivers of broadleaved evergreen species spread into deciduous forests in the southern Swiss Alps’, Regional Environmental Change, 18(2), pp. 425–436. doi: 10.1007/s10113-017-1212-7.

Crous, K. Y., Uddling, J. and De Kauwe, M. G. (2022) ‘Temperature responses of photosynthesis and respiration in evergreen trees from boreal to tropical latitudes’, New Phytologist, 234(2), pp. 353–374. doi: 10.1111/nph.17951.

Davidson, A. M., Jennions, M. and Nicotra, A. B. (2011) ‘Do invasive species show higher phenotypic plasticity than native species and, if so, is it adaptive? A meta-analysis’, Ecology Letters, 14(4), pp. 419–431. doi: 10.1111/j.1461-0248.2011.01596.x.

Didion-Gency, M. et al. (2022) ‘Impact of warmer and drier conditions on tree photosynthetic properties and the role of species interactions’, New Phytologist, 236(2), pp. 547–560. doi: 10.1111/nph.18384.

Didion-Gency, M. et al. (2024) ‘Chronic warming and dry soils limit carbon uptake and growth despite a longer growing season in beech and oak’, Plant Physiology, 194(2), pp. 741–757. doi: 10.1093/plphys/kiad565.

Duan, B. et al. (2013) ‘Plastic responses of Populus yunnanensis and Abies faxoniana to elevated atmospheric CO2 and warming’, Forest Ecology and Management, 296, pp. 33–40. doi: 10.1016/j.foreco.2013.01.032.

Dusenge, M. E., Duarte, A. G. and Way, D. A. (2019) ‘Plant carbon metabolism and climate change: elevated CO2 and temperature impacts on photosynthesis, photorespiration and respiration’, New Phytologist, 221(1), pp. 32–49. doi: 10.1111/nph.15283.

Dusenge, M. E., Madhavji, S. and Way, D. A. (2020) ‘Contrasting acclimation responses to elevated CO2 and warming between an evergreen and a deciduous boreal conifer’, Global Change Biology, 26(6), pp. 3639–3657. doi: 10.1111/gcb.15084.

Fehr, V. et al. (2023) ‘The invasive Chinese windmill palm (Trachycarpus fortunei) impacts forest vegetation and regeneration on the southern slope of the European Alps’, Applied Vegetation Science, (May 2023), pp. 1–31. doi: 10.1111/avsc.12765.

Fehr, V. and Burga, C. A. (2016) ‘Aspects and Causes of Earlier and Current Spread of Trachycarpus fortunei in the Forests of Southern Ticino and’, Palms, 60(3), pp. 125–136. Available at: http://www.zora.uzh.ch/id/eprint/127489/.

Feng, Y. L. and Fu, G. L. (2008) ‘Nitrogen allocation, partitioning and use efficiency in three invasive plant species in comparison with their native congeners’, Biological Invasions, 10(6), pp. 891–902. doi: 10.1007/s10530-008-9240-3.

Finch, D. M. et al. (2021) Effects of Climate Change on Invasive Species, Invasive Species in Forests and Rangelands of the United States. doi: 10.1007/978-3-030-45367-1_17.

Fridley, J. D. (2012) ‘Extended leaf phenology and the autumn niche in deciduous forest invasions’, Nature, 485(7398), pp. 359–362. doi: 10.1038/nature11056.

Fridley, J. D. et al. (2022) ‘A general hypothesis of forest invasions by woody plants based on whole-plant carbon economics’, Journal of Ecology, (April 2022), pp. 4–22. doi: 10.1111/1365-2745.14001.

Funk, J. L., Glenwinkel, L. A. and Sack, L. (2013) ‘Differential Allocation to Photosynthetic and Non-Photosynthetic Nitrogen Fractions among Native and Invasive Species’, PLoS ONE, 8(5). doi: 10.1371/journal.pone.0064502.

Gauthey, A. et al. (2023) ‘Absence of canopy temperature variation despite stomatal adjustment in Pinus sylvestris under multidecadal soil moisture manipulation’, New Phytologist, 240(1), pp. 127–137. doi: 10.1111/nph.19136.

Gelman, A. and Rubin, D. B. (1992) ‘Inference from Iterative Simulation Using Multiple Sequences’, Statistical Science, 7(4), pp. 457–511. doi: 10.2307/2246134.

Gioria, M. et al. (2023) ‘Why Are Invasive Plants Successful?’, Annual Review of Plant Biology, 74, pp. 635–670. doi: 10.1146/annurev-Arplant-070522-071021.

Godoy, O., Valladares, F. and Castro-Díez, P. (2011) ‘Multispecies comparison reveals that invasive and native plants differ in their traits but not in their plasticity’, Functional Ecology, 25(6), pp. 1248–1259. doi: 10.1111/j.1365-2435.2011.01886.x.

Grossiord, C. et al. (2022) ‘Warming may extend tree growing seasons and compensate for reduced carbon uptake during dry periods’, Journal of Ecology, (April), pp. 1–15. doi: 10.1111/1365-2745.13892.

Gunderson, C. A. et al. (2010) ‘Thermal plasticity of photosynthesis: The role of acclimation in forest responses to a warming climate’, Global Change Biology, 16(8), pp. 2272–2286. doi: 10.1111/j.1365-2486.2009.02090.x.

Hartig, F. et al. (2023) ‘General-Purpose MCMC and SMC Samplers and Tools for Bayesian Statistics’. Available at: https://github.com/florianhartig/BayesianTools.

Higgins, S. I. and Richardson, D. M. (2014) ‘Invasive plants have broader physiological niches’, Proceedings of the National Academy of Sciences of the United States of America, 111(29), pp. 10610–10614. doi: 10.1073/pnas.1406075111.

Jones, H. G. (1992) Plants and Microclimate: A Quantitative Approach to Environmental Plant Physiology: Third Edition. Cambridge University Press.

Juillard, T. et al. (2024) ‘Invasive palms have more efficient and prolonged CO2 assimilation compared to native sub-Mediterranean vegetation’, Forest Ecology and Management, 556(January), p. 121743. doi: 10.1016/j.foreco.2024.121743.

Kattge, J. and Knorr, W. (2007) ‘Temperature acclimation in a biochemical model of photosynthesis: A reanalysis of data from 36 species’, Plant, Cell and Environment, 30(9), pp. 1176–1190. doi: 10.1111/j.1365-3040.2007.01690.x.

Kruse, J. et al. (2019) ‘Optimization of photosynthesis and stomatal conductance in the date palm Phoenix dactylifera during acclimation to heat and drought’, New Phytologist, 223(4), pp. 1973–1988. doi: 10.1111/nph.15923.

Kullberg, A. T., Slot, M. and Feeley, K. J. (2023) ‘Thermal optimum of photosynthesis is controlled by stomatal conductance and does not acclimate across an urban thermal gradient in six subtropical tree species’, Plant Cell and Environment, 46(3), pp. 831–849. doi: 10.1111/pce.14533.

Kumarathunge, D. P. et al. (2019) ‘Acclimation and adaptation components of the temperature dependence of plant photosynthesis at the global scale’, New Phytologist, 222(2), pp. 768–784. doi: 10.1111/nph.15668.

Lee, T. D., Reich, P. B. and Bolstad, P. V. (2005) ‘Acclimation of leaf respiration to temperature is rapid and related to specific leaf area, soluble sugars and leaf nitrogen across three temperate deciduous tree species’, Functional Ecology, 19(4), pp. 640–647. doi: 10.1111/j.1365-2435.2005.01023.x.

Leishman, M. R. et al. (2007) ‘Leaf trait relationships of native and invasive plants: Community- and global-scale comparisons’, New Phytologist, 176(3), pp. 635–643. doi: 10.1111/j.1469-8137.2007.02189.x.

Liu, C. et al. (2020) ‘Most invasive species largely conserve their climatic niche’, Proceedings of the National Academy of Sciences of the United States of America, 117(38), pp. 23643– 23651. doi: 10.1073/pnas.2004289117.

Liu, Y. et al. (2017) ‘Do invasive alien plants benefit more from global environmental change than native plants?’, Global Change Biology, 23(8), pp. 3363–3370. doi: 10.1111/gcb.13579.

Marias, D. E., Meinzer, F. C. and Still, C. (2017) ‘Impacts of leaf age and heat stress duration on photosynthetic gas exchange and foliar nonstructural carbohydrates in Coffea arabica’, Ecology and Evolution, 7(4), pp. 1297–1310. doi: 10.1002/ece3.2681.

Monteith, J. L. and Reifsnyder, W. E. (2007) Principles of Environmental Physics: 3rd Edition, Physics Today. Edited by ELSEVIER. doi: 10.1063/1.3128494.

Montesinos, D. (2022) ‘Fast invasives fastly become faster: Invasive plants align largely with the fast side of the plant economics spectrum’, Journal of Ecology, 110(5), pp. 1010–1014. doi: 10.1111/1365-2745.13616.

Niinemets, Ü., Valladares, F. and Ceulemans, R. (2003) ‘Leaf-level phenotypic variability and plasticity of invasive Rhododendron ponticum and non-invasive Ilex aquifolium co-occurring at two contrasting European sites’, Plant, Cell and Environment, 26(6), pp. 941–956. doi: 10.1046/j.1365-3040.2003.01027.x.

O’Sullivan, O. S. et al. (2013) ‘High-resolution temperature responses of leaf respiration in snow gum (Eucalyptus pauciflora) reveal high-temperature limits to respiratory function’, Plant, Cell and Environment, 36(7), pp. 1268–1284. doi: 10.1111/pce.12057.

Pearcy, R. W. et al. (1987) ‘Carbon Gain by Plants in Natural Environments’, BioScience, 37(1), pp. 21–29. doi: 10.2307/1310174.

Petrik, P. et al. (2022) ‘Interannual adjustments in stomatal and leaf morphological traits of European beech (Fagus sylvatica L.) demonstrate its climate change acclimation potential’, Plant Biology, 24(7), pp. 1287–1296. doi: 10.1111/plb.13401.

Pyšek, P. et al. (2020) ‘Scientists’ warning on invasive alien species’, Biological Reviews, 95(6), pp. 1511–1534. doi: 10.1111/brv.12627.

Quan, X., Wang, N. and Wang, C. (2020) ‘Thermal acclimation of leaf dark respiration of Larix gmelinii: A latitudinal transplant experiment’, Science of the Total Environment, 743, p. 140634. doi: 10.1016/j.scitotenv.2020.140634.

Reich, P. B. et al. (1998) ‘Relationships of leaf dark respiration to leaf nitrogen, specific leaf area and leaf life-span: a test across biomes and functional groups’, Oecologia, pp. 471–482.

Reich, P. B. et al. (2016) ‘Boreal and temperate trees show strong acclimation of respiration to warming’, Nature, 531(7596), pp. 633–636. doi: 10.1038/nature17142.

Ripley, B. S. et al. (2020) ‘Invasive grasses of sub-Antarctic Marion Island respond to increasing temperatures at the expense of chilling tolerance’, pp. 765–773. doi: 10.1093/aob/mcz156.

Schindelin, J. et al. (2012) ‘Fiji: An open-source platform for biological-image analysis’, Nature Methods, 9(7), pp. 676–682. doi: 10.1038/nmeth.2019.

Sendall, K. M., Lusk, C. H. and Reich, P. B. (2016) ‘Trade-offs in juvenile growth potential vs. shade tolerance among subtropical rain forest trees on soils of contrasting fertility’, Functional Ecology, 30(6), pp. 845–855. doi: 10.1111/1365-2435.12573.

Slot, M. et al. (2024) ‘The stomatal response to vapor pressure deficit drives the apparent temperature response of photosynthesis in tropical forests’, New Phytologist. doi: 10.1111/nph.19806.

Slot, M. and Kitajima, K. (2015) ‘General patterns of acclimation of leaf respiration to elevated temperatures across biomes and plant types’, Oecologia, 177(3), pp. 885–900. doi: 10.1007/s00442-014-3159-4.

Slot, M. and Winter, K. (2017) ‘Photosynthetic acclimation to warming in tropical forest tree Seedlings’, Journal of Experimental Botany, 68(9), pp. 2275–2284. doi: 10.1093/jxb/erx071.

Smith, N. G. and Dukes, J. S. (2013) ‘Plant respiration and photosynthesis in global-scale models: Incorporating acclimation to temperature and CO2’, Global Change Biology, 19(1), pp. 45–63. doi: 10.1111/j.1365-2486.2012.02797.x.

Smith, N. G. and Dukes, J. S. (2017) ‘Short-term acclimation to warmer temperatures accelerates leaf carbon exchange processes across plant types’, Global Change Biology, 23(11), pp. 4840–4853. doi: 10.1111/gcb.13735.

Sridhar, V. and Elliott, R. L. (2002) ‘On the development of a simple downwelling longwave radiation scheme’, Agricultural and Forest Meteorology, 112(3–4), pp. 237–243. doi: 10.1016/S0168-1923(02)00129-6.

Still, C. J. et al. (2019) ‘When a cuvette is not a canopy: A caution about measuring leaf temperature during gas exchange measurements’, Agricultural and Forest Meteorology, 279(November 2018), p. 107737. doi: 10.1016/j.agrformet.2019.107737.

Turbelin, A. and Catford, J. A. (2021) Invasive plants and climate change, Climate Change: Observed Impacts on Planet Earth, Third Edition. Elsevier B.V. doi: 10.1016/B978-0-12-821575-3.00025-6.

Verlinden, M. and Nijs, I. (2010) ‘Alien plant species favoured over congeneric natives under experimental climate warming in temperate Belgian climate’, pp. 2777–2787. doi: 10.1007/s10530-009-9683-1.

Vico, G. et al. (2019) ‘Can leaf net photosynthesis acclimate to rising and more variable temperatures?’, Plant Cell and Environment, 42(6), pp. 1913–1928. doi: 10.1111/pce.13525.

Walther, G. R. et al. (2007) ‘Palms tracking climate change’, Global Ecology and Biogeography, 16(6), pp. 801–809. doi: 10.1111/j.1466-8238.2007.00328.x.

Walther, G. R. et al. (2009) ‘Alien species in a warmer world: risks and opportunities’, Trends in Ecology and Evolution, 24(12), pp. 686–693. doi: 10.1016/j.tree.2009.06.008.

Wang, H. et al. (2020) ‘Acclimation of leaf respiration consistent with optimal photosynthetic capacity’, Global Change Biology, 26(4), pp. 2573–2583. doi: 10.1111/gcb.14980.

Way, D. A. and Oren, R. (2010) ‘Differential responses to changes in growth temperature between trees from different functional groups and biomes: a review and synthesis of data’, Tree Physiology, 30(6), pp. 669–688. doi: 10.1093/treephys/tpq015.

Way, D. A. and Yamori, W. (2014) ‘Thermal acclimation of photosynthesis : on the importance of adjusting our definitions and accounting for thermal acclimation of respiration’, pp. 89–100. doi: 10.1007/s11120-013-9873-7.

van de Weg, M. J., Fetcher, N. and Shaver, G. (2013) ‘Response of dark respiration to temperature in Eriophorum vaginatum from a 30-year-old transplant experiment in Alaska’, Plant Ecology and Diversity, 6(3–4), pp. 377–381. doi: 10.1080/17550874.2012.729618.

Xu, C. Y., Schuster, W. S. F. and Griffin, K. L. (2007) ‘Seasonal variation of temperature response of respiration in invasive Berberis thunbergii (Japanese barberry) and two co-occurring native understory shrubs in a northeastern US deciduous forest’, Oecologia, 153(4), pp. 809–819. doi: 10.1007/s00442-007-0790-3.

Yamori, W., Hikosaka, K. and Way, D. A. (2014) ‘Temperature response of photosynthesis in C3, C4, and CAM plants: Temperature acclimation and temperature adaptation’, Photosynthesis Research, 119(1–2), pp. 101–117. doi: 10.1007/s11120-013-9874-6.

Yu, L. et al. (2019) ‘Elevated temperature differently affects growth, photosynthetic capacity, nutrient absorption and leaf ultrastructure of Abies faxoniana and Picea purpurea under intra- And interspecific competition’, Tree Physiology, 39(8), pp. 1342–1357. doi: 10.1093/treephys/tpz044.

Zhu, L. et al. (2020) ‘Acclimation of leaf respiration temperature responses across thermally contrasting biomes’, New Phytologist. doi: 10.1111/nph.16929.

