## Supplementary material for "Thermal acclimation fails to confer a carbon budget advantage to invasive species over natives"

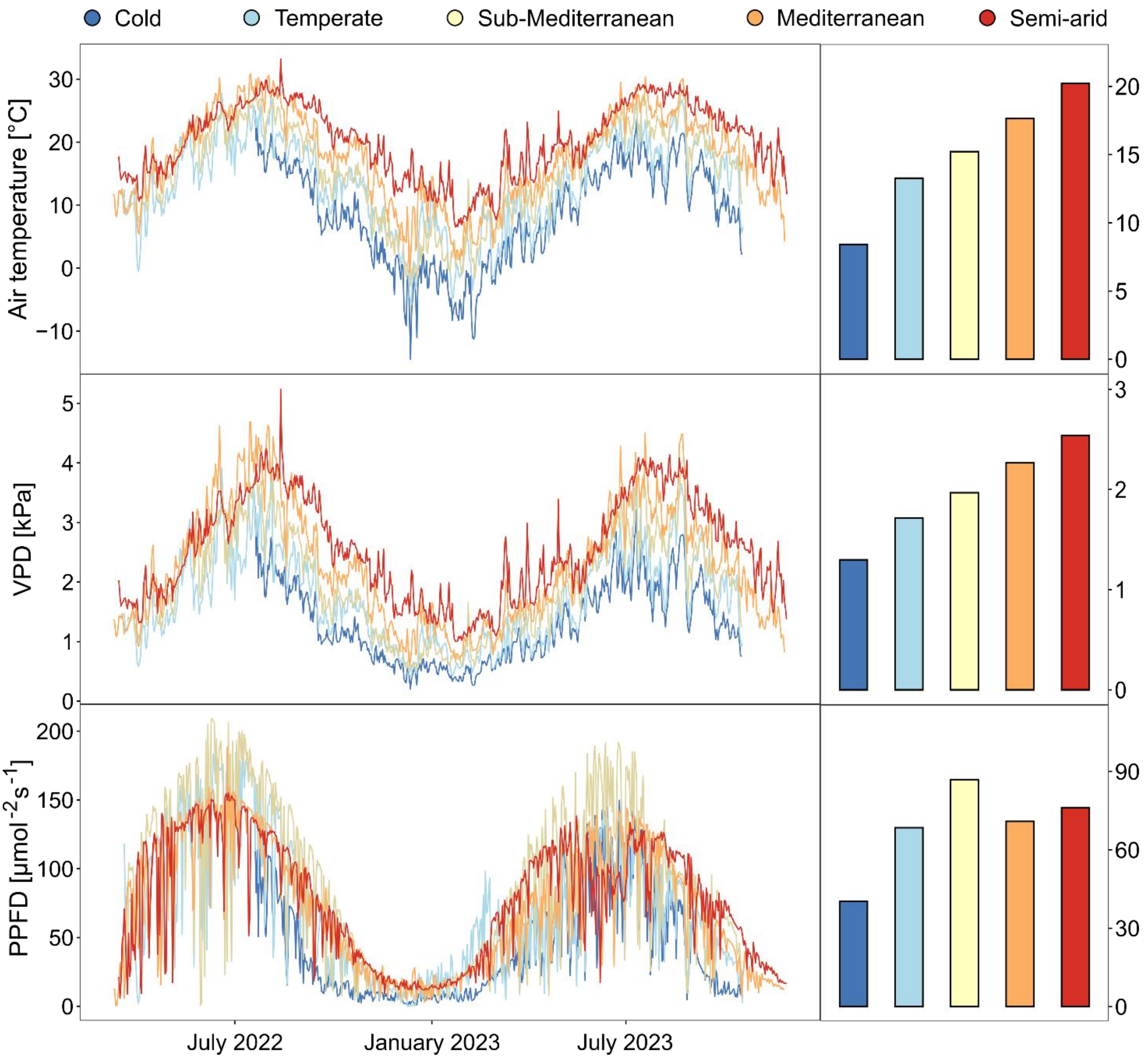


**Fig. S1:** Mean daily air temperature (top), vapor pressure deficit (VPD; middle), and photosynthetically active flux density (PPFD; bottom) within the shading infrastructures of the five experimental sites from March 2022 to October 2023. The yearly mean values of the three variables are presented on the right panel by sites.

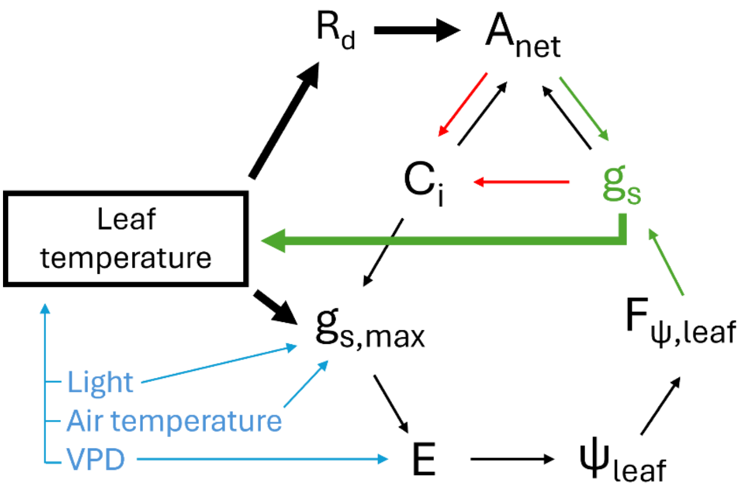


**Fig. S2**: Simplified representation of the SPAC model. The optimization procedure to find a stable stomatal conductance ($g_s$, thin arrows) is nested within the optimization procedure to find a stable leaf temperature (thick arrows). Colors are used to show different steps of the process. (1) Leaf temperature is calculated from environmental drivers (in blue) with an energy balance module. (2) Initial values of intercellular $CO_2$ concentration ($C_i$) and $g_s$ are used to initiate the optimization procedure on $g_s$. (3) Iteratively, $g_s$ is optimized from the net photosynthesis rate ($A_{net}$) and a correction factor ($F_{\psi,leaf}$) (in thin green) after computing the maximal stomatal conductance ($g_{s,max}$), evaporation (E), and leaf water potential ($\psi_{leaf}$). The iterative process stops when the new $C_i$, calculated with the optimized $g_s$ (in red), matches the Ci of the previous iteration. (5) Optimized $g_s$ is used to calculate a new leaf temperature compared with the initial leaf temperature (thick green arrow). New iterations on both loops are run as long as values differ.

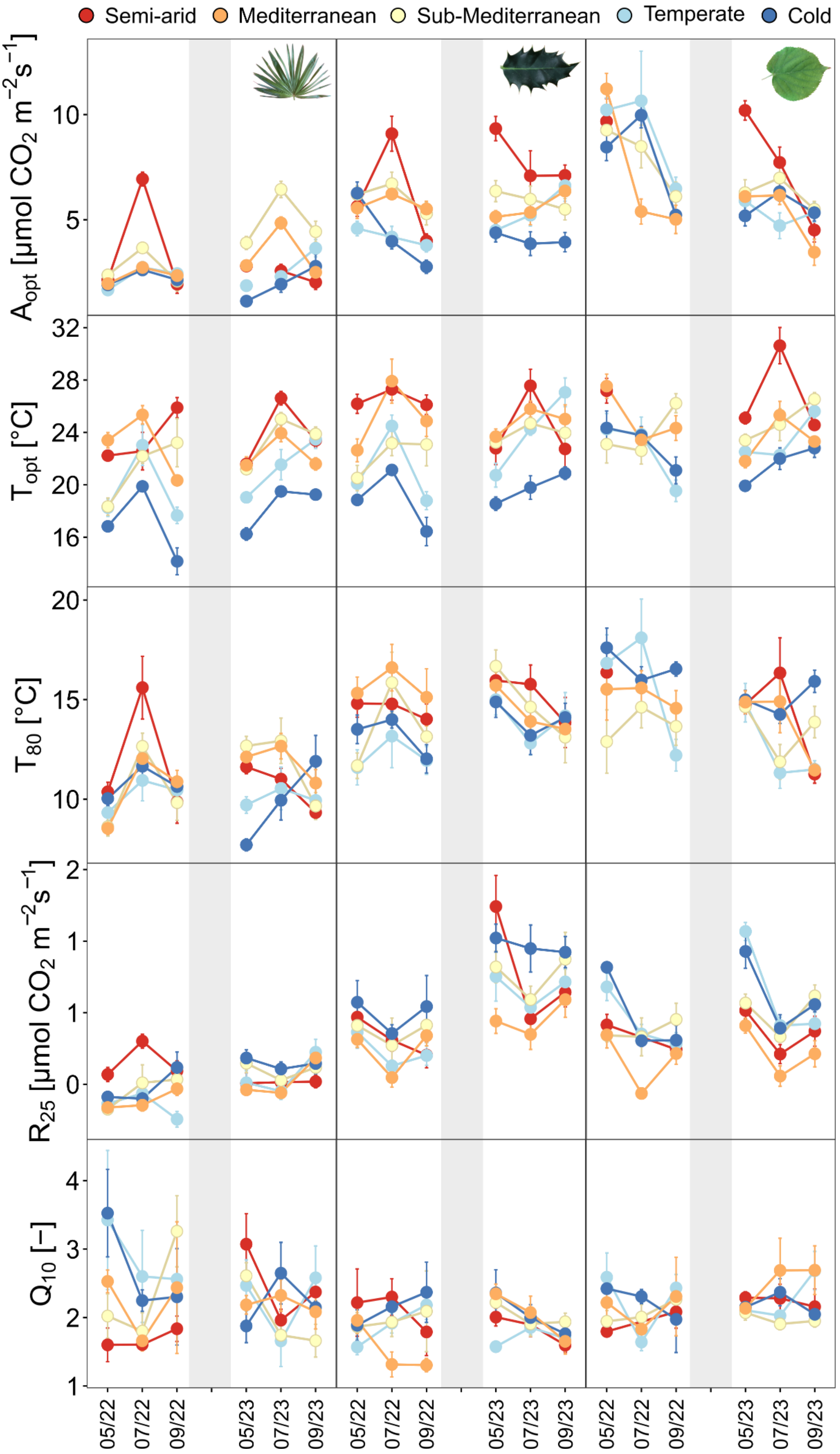
Semi-arid
Mediterranean
Sub-Mediterranean
Temperate
Cold
$A_{opt}$ [µmol $CO_2$ $m^{-2}s^{-1}$]
$T_{opt}$ [°C]
$T_{80}$ [°C]
$R_{25}$ [µmol $CO_2$ $m^{-2}s^{-1}$]
$Q_{10}$ [–]
05/22
07/22
09/22
05/23
07/23
09/23

**Fig. S3**: Optimal net assimilation ($A_{opt}$), optimal temperature ($T_{opt}$), thermal breathing ($T_{80}$), respiration at 25°C ($R_{25}$), and respiration yield per 10°C increase ($Q_{10}$) (means ± s.e., n = 4-10 individuals per species) of all species during the six measurement campaigns in 2022-2023. Significant differences between species (Tukey's HSD post hoc test, alpha = 0.05) at each campaign are indicated with different letters in Table S5.

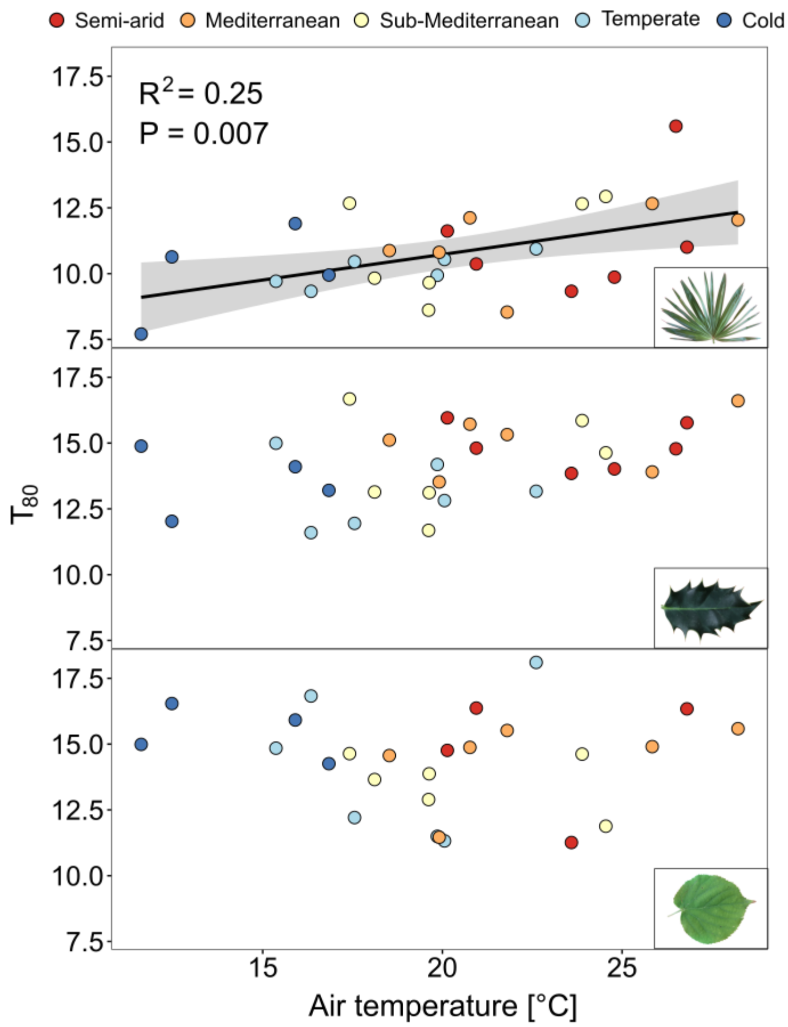


**Fig. S4:** Relationships between $T_{80}$ (n = 4-10 individuals per species) averaged by campaigns and air temperature of the two weeks preceding the measurements for *T. fortunei*, *I. aquifolium*, and *T. cordata*. Colors represent climates from blue to red, going from the coldest to the warmest. The regression lines were fitted with a linear model when significant.

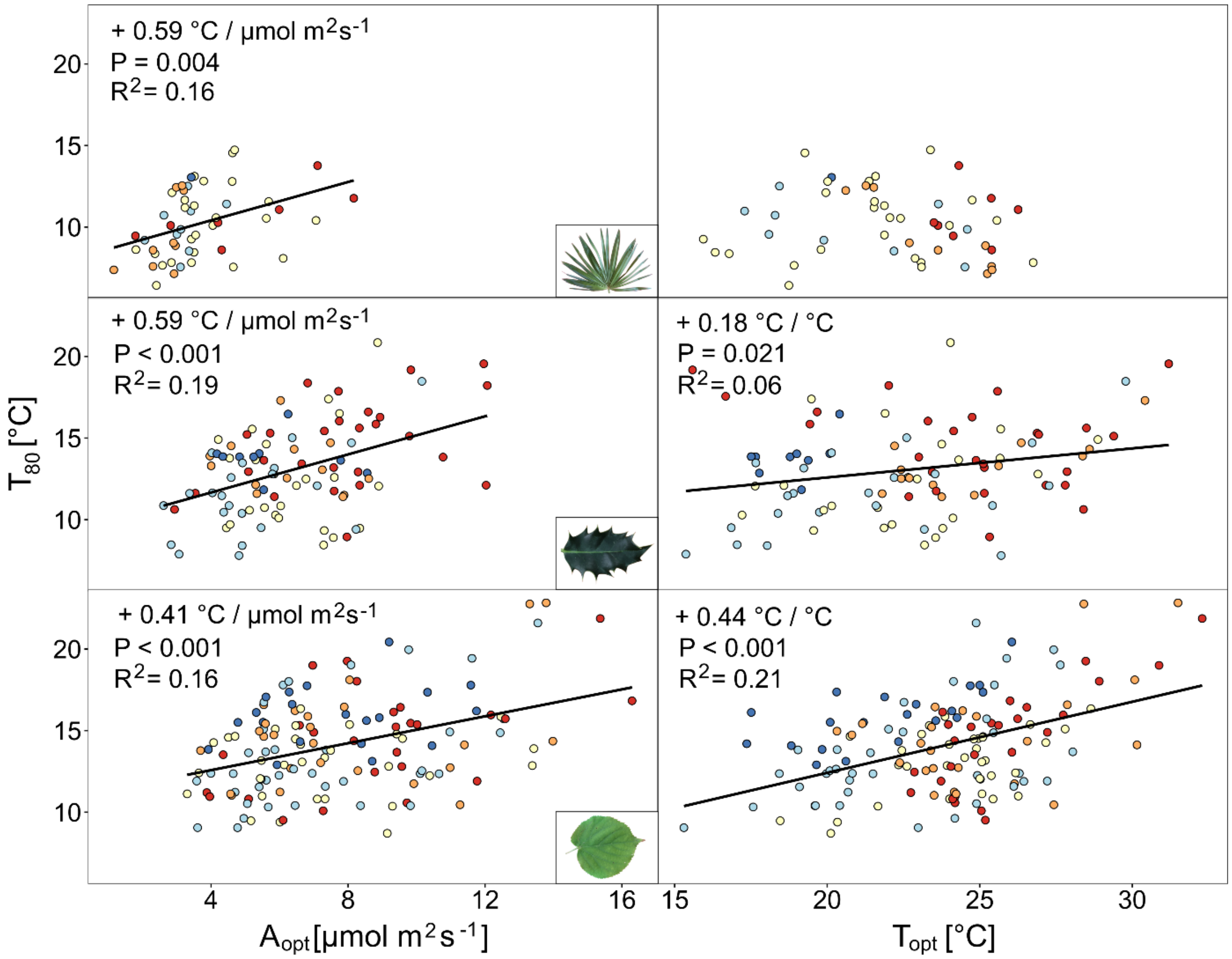


**Fig. S5:** Thermal breath ($T_{80}$) in function of the assimilation at the optimal temperature ($A_{opt}$) and optimal temperature ($T_{opt}$) for *T. fortunei*, *I. aquifolium*, and *T. cordata* during all campaigns in 2022 and 2023. Colors represent sites from blue to red, going from the coldest to the warmest. The regression lines were fitted with a linear model when significant.

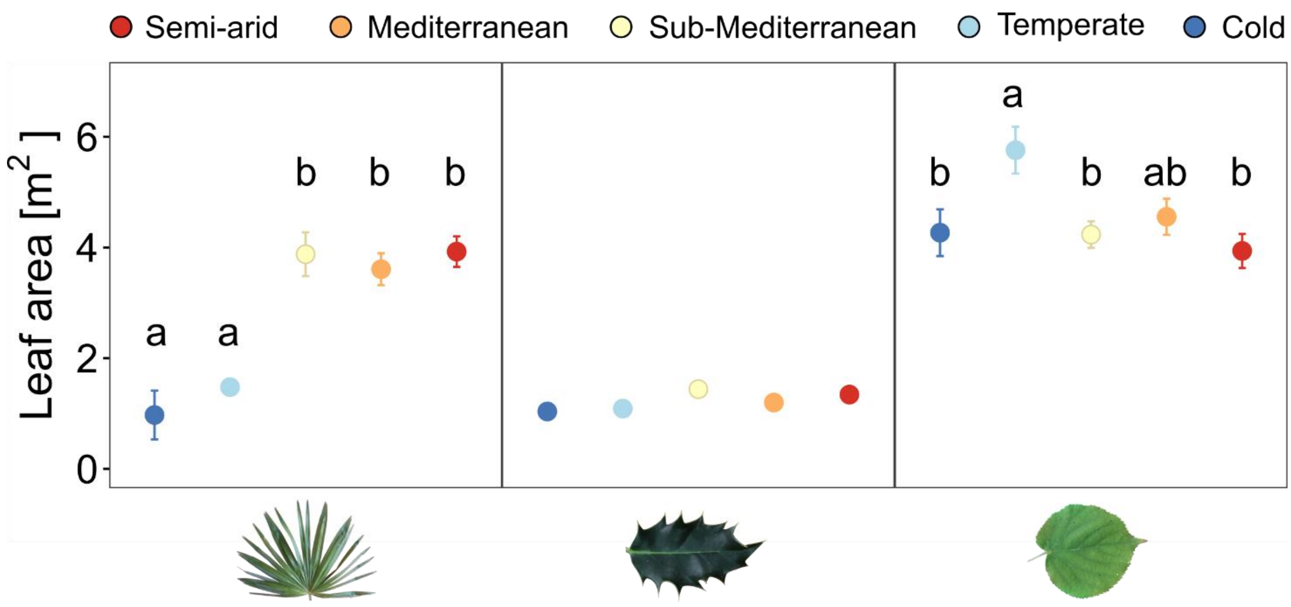


33 **Fig. S6**: Leaf area (means +/- s.e., n = 4-10 individuals per species) of all species and sites

34 in September 2023.

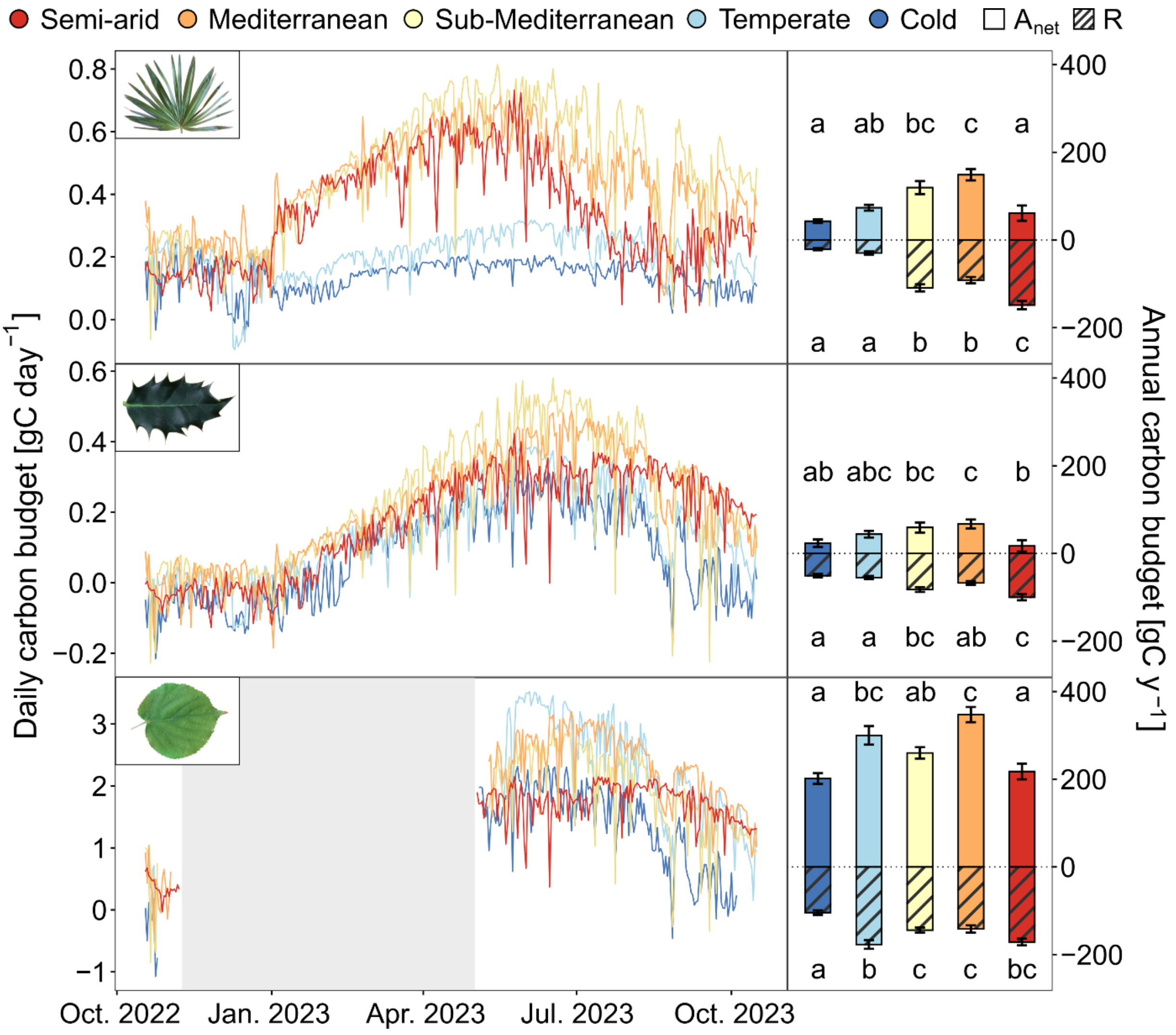


**Fig. S7:** Daily mean net C uptake of *T. fortunei*, *I. aquifolium*, and *T. cordata* at the five experimental sites from 15th October 2022 to 15th October 2023, multiplied by the leaf area. The right panels show the yearly C uptake at each site. Net assimilation corresponds to the plain bars, whereas respiration bars are dashed in black. Error bars indicate the uncertainty of $J_{max,25}$, $V_{Cmax,25}$, $R_{25}$, and $Q_{10}$ modeled (n = 37-57).

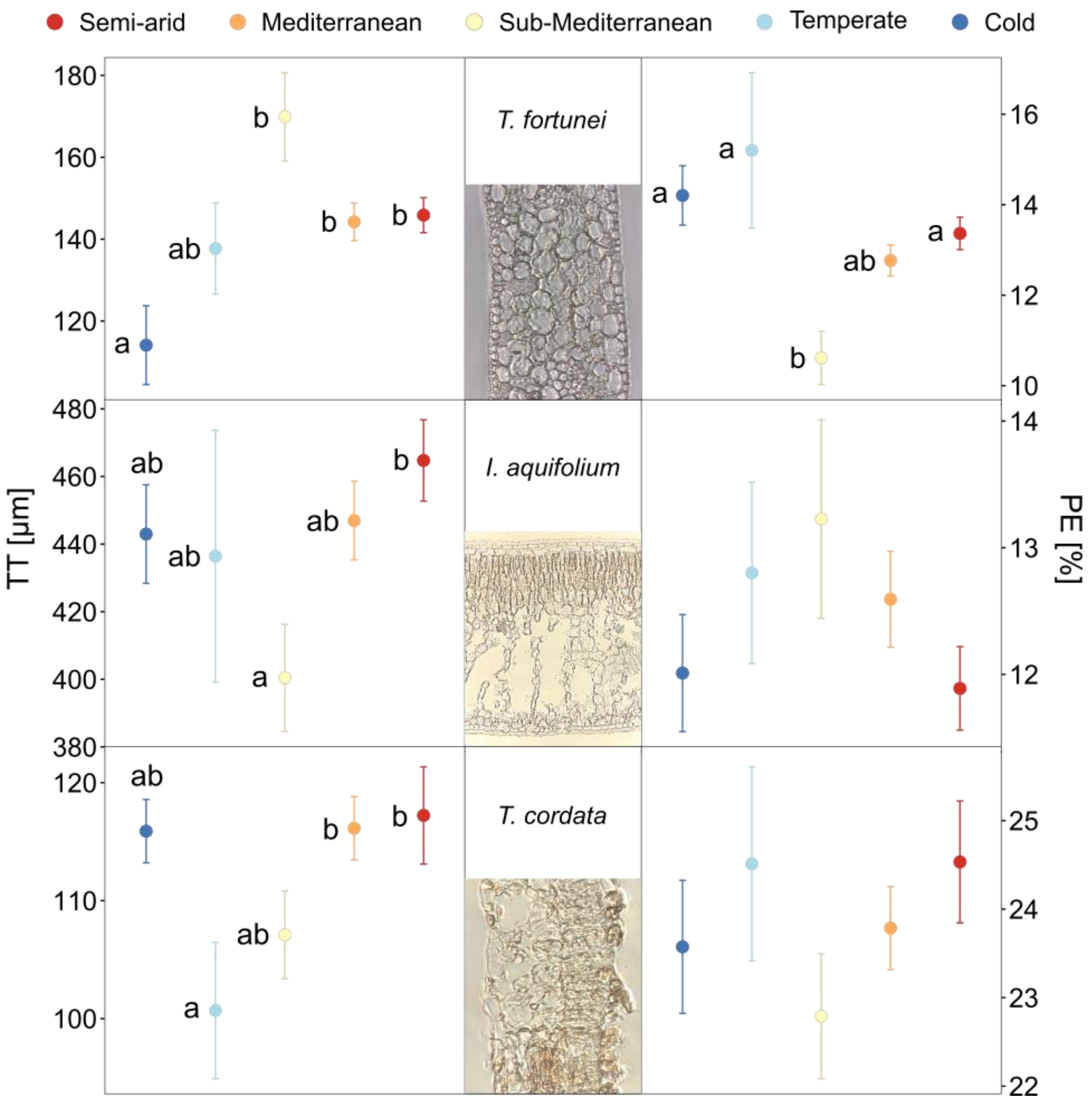


**Fig. S8:** Leaf total thickness (TT) and percentage of the epidermis (PE) relative to the total leaf thickness (n = 3-10 individuals per species) for *T. fortunei*, *I. aquifolium*, and *T. cordata* growing in the five experimental sites. Samples were collected in September 2023 after a year and a half of acclimation in the different sites. Significant differences between the treatments are indicated with letters (Tukey's HSD post-hoc test, alpha = 0.05).

**Table S1**: List of the input parameters used in the SPAC model.

| Variable | Symbol in LPM R package | Unit | Value | | | Source |
|---|---|---|---|---|---|---|
| *Environmental parameters* | | | | | | |
| Air temperature | Ta | °C | | | | Measured |
| Air humidity | RH | % | | | | Measured |
| Light availability | L | PPFD | | | | Measured |
| Wind speed | WS | $m\ s^{-1}$ | | 2 | | - |
| Ambient CO2 | Ca | $\mu mol\ mol^{-1}$ | | 400 | | - |
| *Photosynthetic parameters* | | | *T. fortunei* | *I .aquifolium* | *T .cordata* | |
| Max. catalytic activity of Rubisco at 25 °C | vcmax | $\mu mol\ m^{-2}\ s^{-1}$ | 25.97 | 33.48 | 32.18 | Measured |
| Max. ratio of electron transport at 25 °C | jmax | $\mu mol\ m^{-2}\ s^{-1}$ | 62.72 | 84.11 | 61.90 | Measured |
| Respiration at 25 °C | Rd_25 | $\mu mol\ m^{-2}\ s^{-1}$ | 0.41 | 0.85 | 0.75 | Measured |
| Respiration yield | $Q_{10}$ | - | 2.297838 | 1.934928 | 2.182466 | Measured |
| Leaf area index | LAI | $m^2\ m^{-2}$ | 1.907561 | 0.709583 | 3.173085 | Measured |
| Fraction of sunlit leaves | f_leaf_sun | - | | 1 | | - |
| Reference leaf water potential | Psi_f | kPa | -462 | -445 | -470 | Calibrated |
| Shape factor | s_f | $kPa^{-1}$ | 0.0061 | 0.0061 | 0.0980 | Calibrated |
| Minimum stomatal conductance | $g_{min}$ | $\mu mol\ m^{-2}\ s^{-1}$ | 0.0038 | 0.0028 | 0.0060 | Measured |
| ProportionS.-A.ty factor | m | - | 0.02 | 0.17 | 0.3 | Calibrated |
| Degree of curvature PAR / $J_{max}$ | gamma | $\mu mol\ m^{-2}$ | | 0.9 | | Buckley et al. (2014)<br>Garcia-Tejera (2023) |
| *Hydrological parameters* | | | | | | |
| Soil water potential | Psi_soil | kPa | | -300 | | Garcia-Tejera (2023) |
| Root resistance | R_root | $kPa\ m^2\ s\ kg^{-1}$ | | 10000 | | Grossiord (2022) |
| Xylem resistance | R_xylem | $kPa\ m^2\ s\ kg^{-1}$ | 0.0625 | 0.0766 | 0.0499 | Grossiord (2022) |
| Soil resistance | R_soil | kPa | | 100000 | | Garcia-Tejera (2023) |
| *Leaf temperature* | | | | | | |
| Leaf angle from horizontal | i | ° | | 30 | | Leaf Energy Balance function<br>(http://landflux.org/Tools.php) |
| Absorptance to SWR | $a_{SWR}$ | % | | 0.5 | | |
| Emissivity | em | - | | 0.97 | | |
| Characteristic dimension | d | mm | 320 | 40 | 60 | Measured |
| Shape of the leaf | shape_index | Cat. | 1 | 2 | 1 | Measured |
| *Photosynthetic parameters for thermal acclimation* | | | | | | |
| Activation energy $J_{max}$ | Ha | $kJ\ mol^{-1}$ | | 51.5 | | |
| Activation energy $V_{Cmax}$ | | | | 79.4 | | |
| Deactivation energy | Hd | $kJ\ mol^{-1}$ | | 200 | | |
| Entropy factor $J_{max}$ | Delta_S | $°C^{-1}$ | | 662.3 | | Kumarathunge et al. (2019) |
| Entropy factor $V_{Cmax}$ | | | | 647.9 | | |
| *Respiration parameters for thermal acclimation* | | | | | | |
| Slope for $R_{25}$ acclimation | | $\mu mol\ m^{-2}\ s^{-1}$ / °C | -0.0037 | -0.0293 | -0.0317 | Measured |
| Intercept for $R_{25}$ acclimation | | $\mu mol\ m^{-2}\ s^{-1}$ | 0.50 | 1.42 | 1.36 | Measured |

**Table S2**: Variables, equations, and descriptions of the SPAC model

| **Var.** | **Symbol** | **Unit** | **Equation** | **Definition of var. in the equation** |
|---|---|---|---|---|
| Optimization process over $T_L$ | | | | |
| Dark respiration | R | μmol $m^2$ $s^{-1}$ | $R_{25}$ * $Q_{10}^{TL\text{-}25\ /\ 10}$ | $Q_{10}$ = Respiration yield; $T_L$ = leaf temperature |
| Max. catalytic activity of Rubisco at 25 °C | $V_{Cmax}$ | μmol $m^{-2}$ $s^{-1}$ | Equation (5) in the main manuscript | |
| Max. ratio of electron transport at 25 °C | $J_{max}$ | μmol $m^{-2}$ $s^{-1}$ | Equation (5) in the main manuscript | |
| Activation energy for $V_{Cmax}$ | $H_a$ | kJ $mol^{-1}$ | 39.7 + 1.14 $T_{growth}$ | $T_{growth}$ = mean air temperature of the two previous weeks |
| Activation energy for $J_{max}$ | $H_a$ | kJ $mol^{-1}$ | 27.2 + 0.26 $T_{growth}$ | |
| Entropy factor for $V_{Cmax}$ | ΔS | J $mol^{-1}$ $K^{-1}$ | 645.1 – 0.38 $T_{growth}$ | |
| Entropy factor for $J_{max}$ | ΔS | J $mol^{-1}$ $K^{-1}$ | 653.9 – 0.85 $T_{growth}$ | |
| Optimization process over $g_s$ | | | | |
| Maximal stomatal conductance | $g_{s,max}$ | μmol $m^{-2}$ $s^{-1}$ | Minimum of [B ($C_i$ - Γ) - R (E$C_i$ + D)] / [((E$C_i$ + D)($C_i$ - $C_a$)] for $VC_{max}$ or J, see Garcia *et al.*, (2017) for details | $C_i$ = Internal $CO_2$ concentration [μmol $m^{-1}$]; $C_a$ = ambient $CO_2$ concentration [μmol $m^{-1}$]; Γ = $CO_2$ compensation point [μmol $CO_2$ $m^{-1}$]; B = $VC_{max}$ or J (calculated from Jmax, quantum efficiency, and APAR) D = 210.1 or 0.8 (combination of Michaelis–Menten coefficients and Γ) E = 1 or 4 |
| Gross assimilation | $A_{gross}$ | μmol $m^{-2}$ $s^{-1}$ | $g_s$ ($C_a$- $C_i$) + R | $C_i$ = ; $C_a$ = ambient $CO_2$ concentration [μmol $m^{-1}$] |
| Transpiration | TR | Kg $s^{-1}$ $m^{-2}$ | $g_{s,max}$ LAI VPD / P | LAI = Leaf area index [$m^2$ $m^{-2}$; VPD = water vapor pressure deficit of the air [kPa]; P = atmospheric pressure [Pa] |
| Leaf water potential | ψleaf | kPa | ($ψ_{soil}$ / ($R_{soil}$ + $R_{root}$)) / (1 / ($R_{soil}$ + $R_{root}$)) – TR * [($R_{soil}$ + $R_{root}$ + $R_{xylem}$) / $f_{leaf\ sun}$] | $Ψ_{soil}$ = ; $R_{soil}$, $R_{root}$, $R_{xylem}$ = soil, root, and xylem hydraulic resistance [s m kPa $kg^{-1}$]; $f_{leaf\ sun}$ = fraction of the LAI illuminated |
| Modifying factor of leaf water potential | $f_{ψ,leaf}$ | - | (1+ $e^{sf\ ψf}$) / (1+ $e^{sf\ (ψf\ –\ ψleaf)}$) | Sf = Shape factor [$kPa^{-1}$]; ψf = Reference leaf water potential [kPa] |
| Stomatal conductance | $g_s$ | μmol $m^{-2}$ $s^{-1}$ | $g_{min}$ + [$f_{ψ,leaf}$ m $A_{gross}$ / ($C_i$ - Γ)] | $g_{min}$ = minimal stomatal conductance [μmol $m^{-2}$ $s^{-1}$] ; m = proportionS.-A.ty factor between photosynthesis and stomatal conductance [-]; $C_i$ = Internal $CO_2$ concentration [μmol $m^{-1}$]; Γ = $CO_2$ compensation point [μmol $CO_2$ $m^{-1}$] |

**Table S3**: Variables, equations, and descriptions of the leaf energy balance model

| Var. | Symbol | Unit | Equation | Definition of var. in the equation |
|---|---|---|---|---|
| $e_{sat}$ | Saturation vapor pressure | kPa | $a\ e^{b\ Tair\ /\ (Tair\ +\ z)}$ | a = 0.61121 [kPa]<br>b = 17.502 [-]<br>z = 240.97 [°C]<br>Tair = air temperature [°C] |
| $e_a$ | Water vapor pressure of the air | kPa | $e_{sat}$ (RH / 100) | RH = relative humidity [%] |
| s | Slope of $e_{sat}$ / T curve | kPa $^oC^{-1}$ | $e_{sat}\ b\ z/((T_{air} + z)^2)$ | b = 17.502 [-]<br>z = 240.97 [°C]<br>Tair = air temperature [°C] |
| VPD | Water vapor pressure deficit of the air | kPa | $e_{sat} - e_a$ | |
| $SWR_{abs}$ | Absorbed short-wave radiation | W $m^{-2}$ | $a_{SWR}$ cos(i) SWR | $a_{SWR}$ = absorptance to SWR [%]<br>i = inclination of the leaf from horizontal [°]<br>SWR = short-wave radiation [W $m^{-2}$] |
| $LWR_{in}$ | Incoming long-wave radiation | W $m^{-2}$ | $1.31(10\ e_a / T_{air})^{(1/7)}$ SB $(T_{air} + 273.15)^4$ | Tair = air temperature<br>SB = Stefan-Boltzman constant = 5.67e-8 [W $m^{-2}$ $K^{-4}$] |
| $LWR_{out,i}$ | Isothermal outgoing long-wave radiation | W $m^{-2}$ | em SB $(T_{air} + 273.15)^4$ | em = emissivity = 0.97 [-]<br>SB = Stefan-Boltzman constant = 5.67e-8 [W $m^{-2}$ $K^{-4}$] |
| Rni | Isothermal net radiation | W $m^{-2}$ | $SWR_{abs} + LWR_{in} - LWR_{out,i}$ | |
| $r_r$ | Radiative resistance | s $m^{-1}$ | P $C_p$ / (4 em SB $(Tair + 273.15)^3$) | P = density of air = 1.292 [kg $m^{-3}$]<br>Cp = heat capacity of dry air = 1010 [J $kg^{-1}$ $K^{-1}$]<br>em = emissivity = 0.97 [-]<br>SB = Stefan-Boltzman constant = $5.67e^{-8}$ W $m^{-2}$ $K^{-4}$ |
| $r_{bl}$ | Leaf boundary-layer resistance | s $m^{-1}$ | $1 / (g_x (WS^{jx} / d^{1-jx})$ | Gx [m] & Jx [-] = coefficients depending on leaf shape (flat = 0.00662 & 0.5; cylinder = 0.00403 & 0.6; sphere = 0.00571 & 0.6)<br>WS = wind speed [m $s^{-1}$]<br>d = characteristic dimension [mm] |
| $r_{blr}$ | Boundary-layer + radiative resistance | s $m^{-1}$ | $1 / (r_{bl}^{-1} + r_r^{-1})$ | |
| $y_m$ | Modified psychrometric constant | kPa $K^{-1}$ | $y\ (r_{st} / r_{blr})$ | y = psychrometric constant = 0.066 [kPa $K^{-1}$] |
| $T_{leaf}$ | Leaf temperature | $^oC$ | $T_{air} + (y_m\ R_{ni}\ r_{blr} / (p\ C_p) - VPD) / (s + y_m)$ | P = density of air = 1.292 [kg $m^{-3}$]<br>Cp = heat capacity of dry air = 1010 [J $kg^{-1}$ $K^{-1}$] |

**Table S4**: Results of the two-way ANOVA testing the effects of climate and species on the leaf-level optimal assimilation ($A_{opt}$), temperature at the optimal assimilation ($T_{opt}$), thermal breath of photosynthesis ($T_{80}$), respiration rate at 25°C ($R_{25}$), and respiration yield ($Q_{10}$). Significant effects (*P* < 0.05) are shown in bold.

| **Variable** | **Fixed effect** | **Df** | **Sum Sq** | **Mean Sq** | **F-value** | **P-values** |
|---|---|---|---|---|---|---|
| $A_{opt}$ | Climate | 4 | 192.2 | 48 | 11.7 | **< 0.001** |
| | Species | 2 | 1972.7 | 986.4 | 239.3 | **< 0.001** |
| | Climate:Species | 8 | 111 | 13.9 | 3.4 | **< 0.001** |
| | Residuals | 652 | 2687 | 4.1 | | |
| $T_{opt}$ | Climate | 4 | 1991 | 497.7 | 59.9 | **< 0.001** |
| | Species | 2 | 972 | 485.9 | 58.5 | **< 0.001** |
| | Climate:Species | 8 | 158 | 19.8 | 2.4 | **0.0158** |
| | Residuals | 652 | 5418 | 8.3 | | |
| $T_{80}$ | Climate | 4 | 97 | 24.1 | 3.3 | **0.0112** |
| | Species | 2 | 1859 | 929.4 | 126.4 | **< 0.001** |
| | Climate:Species | 8 | 157 | 19.7 | 2.7 | **0.0068** |
| | Residuals | 652 | 4795 | 7.4 | | |
| $R_{25}$ | Climate | 4 | 4.4 | 1.1 | 17.3 | **< 0.001** |
| | Species | 2 | 23.9 | 11.9 | 186.9 | **< 0.001** |
| | Climate:Species | 8 | 2.0 | 0.3 | 3.9 | **< 0.001** |
| | Residuals | 682 | 43.6 | 0.1 | | |
| $Q_{10}$ | Climate | 4 | 128 | 31.9 | 2.6 | **0.0352** |
| | Species | 2 | 141 | 70.5 | 5.7 | **0.0034** |
| | Climate:Species | 8 | 411 | 51.4 | 4.2 | **< 0.001** |
| | Residuals | 682 | 8378 | 12.3 | | |

**Table S5**: Tukey's HSD post hoc test of optimal net assimilation ($A_{opt}$), optimal temperature ($T_{opt}$), thermal breathing ($T_{80}$), dark respiration at 25°C ($R_{25}$), and respiration yield per 10°C increase ($Q_{10}$) (means ± s.e., n = 4-10 individuals per species) of all species and campaigns. Significant differences between sites for each species and campaign are indicated with different letters (*P* < 0.05).

| Campaign | Species | Climate | $A_{opt}$ | $T_{opt}$ | $T_{80}$ | $R_{25}$ | $Q_{10}$ | Campaign | Species | Climate | $A_{opt}$ | $T_{opt}$ | $T_{80}$ | $R_{25}$ | $Q_{10}$ |
|---|---|---|---|---|---|---|---|---|---|---|---|---|---|---|---|
| 2023 May | *T. fortunei* | S.-A. | - | b | b | b | - | 2023 May | *T. fortunei* | S.-A. | a | b | b | b | - |
| | | Med. | - | b | a | a | - | | | Med. | a | b | b | - | b |
| | | Sub-Med. | - | a | a | a | - | | | Sub-Med. | c | b | b | - | ab |
| | | Temp. | - | a | ab | a | - | | | Temp. | ab | a | a | - | a |
| | | Cold | - | a | ab | ab | - | | | Cold | b | c | c | - | ab |
| | *I. aquifolium* | S.-A. | - | c | b | - | - | | *I. aquifolium* | S.-A. | c | a | - | - | - |
| | | Med. | - | a | b | - | - | | | Med. | ab | a | - | b | - |
| | | Sub-Med. | - | ab | a | - | - | | | Sub-Med. | b | a | - | ab | - |
| | | Temp. | - | ab | a | - | - | | | Temp. | a | ab | - | b | - |
| | | Cold | - | b | ab | - | - | | | Cold | a | b | - | ab | - |
| | *T. cordata* | S.-A. | - | - | - | ac | - | | *T. cordata* | S.-A. | b | b | - | a | - |
| | | Med. | - | - | - | c | - | | | Med. | a | ac | - | a | - |
| | | Sub-Med. | - | - | - | c | - | | | Sub-Med. | a | ab | - | a | - |
| | | Temp. | - | - | - | ab | - | | | Temp. | a | a | - | b | - |
| | | Cold | - | - | - | b | - | | | Cold | a | c | - | b | - |
| 2023 July | *T. fortunei* | S.-A. | b | ab | b | b | - | 2023 July | *T. fortunei* | S.-A. | a | d | - | b | - |
| | | Med. | a | a | ab | a | - | | | Med. | c | ac | - | - | - |
| | | Sub-Med. | a | ab | ab | ab | - | | | Sub-Med. | b | cd | - | - | - |
| | | Temp. | a | ab | a | a | - | | | Temp. | a | ab | - | - | - |
| | | Cold | a | b | ab | a | - | | | Cold | a | b | - | - | - |
| | *I. aquifolium* | S.-A. | c | b | - | - | b | | *I. aquifolium* | S.-A. | a | b | - | ab | - |
| | | Med. | ab | b | - | - | a | | | Med. | ab | ab | - | b | - |
| | | Sub-Med. | b | a | - | - | ab | | | Sub-Med. | ab | ab | - | ab | - |
| | | Temp. | a | ab | - | - | ab | | | Temp. | ab | a | - | a | - |
| | | Cold | a | a | - | - | b | | | Cold | b | c | - | ab | - |
| | *T. cordata* | S.-A. | | | | | | | *T. cordata* | S.-A. | b | b | b | a | - |
| | | Med. | b | - | - | b | ab | | | Med. | ab | a | ab | a | - |
| | | Sub-Med. | ab | - | - | a | ab | | | Sub-Med. | ab | a | a | a | - |
| | | Temp. | a | - | - | a | a | | | Temp. | a | a | a | ab | - |
| | | Cold | a | - | - | a | b | | | Cold | ab | a | ab | ab | - |
| 2023 Sept | *T. fortunei* | S.-A. | - | c | - | - | - | 2023 Sept | *T. fortunei* | S.-A. | b | ab | - | b | - |
| | | Med. | - | ab | - | b | - | | | Med. | b | bc | - | - | - |
| | | Sub-Med. | - | bc | - | a | - | | | Sub-Med. | a | a | - | - | - |
| | | Temp. | - | a | - | b | - | | | Temp. | ab | ab | - | - | - |
| | | Cold | - | d | - | b | - | | | Cold | ab | c | - | - | - |
| | *I. aquifolium* | S.-A. | ab | b | - | - | - | | *I. aquifolium* | S.-A. | a | b | - | - | ab |
| | | Med. | b | b | - | - | - | | | Med. | ab | ab | - | - | - |
| | | Sub-Med. | b | b | - | - | - | | | Sub-Med. | ab | ab | - | - | - |
| | | Temp. | a | a | - | - | - | | | Temp. | a | a | - | - | - |
| | | Cold | a | a | - | - | - | | | Cold | b | b | - | - | - |
| | *T. cordata* | S.-A. | | | | | | | *T. cordata* | S.-A. | - | abc | a | - | - |
| | | Med. | - | bc | ab | - | - | | | Med. | - | bc | ab | ab | - |
| | | Sub-Med. | - | b | ab | - | - | | | Sub-Med. | - | a | bc | ab | - |
| | | Temp. | - | a | a | - | - | | | Temp. | - | ab | a | ab | - |
| | | Cold | - | ac | b | - | - | | | Cold | - | c | c | a | - |

**Table S6**: Squared-R ($R^2$), percentage of bias (%Bias), and Nash-Sutcliff-efficiency (NSE) between measured and modeled data with and without acclimation for the five sites and the three species.

| | **Acclimation in model** | | | **No acclimation in model** | | |
|---|---|---|---|---|---|---|
| Species | $R^2$ | %Bias | NSE | $R^2$ | %Bias | NSE |
| *T. fortunei* | 0.17 | -0.69 | 0.11 | 0.15 | -0.83 | 0.09 |
| *I. aquifolium* | 0.61 | 1.41 | 0.47 | 0.61 | 1.54 | 0.42 |
| *T. cordata* | 0.66 | 1.72 | 0.62 | 0.69 | 2.57 | 0.62 |
| ALL | 0.61 | 0.91 | 0.54 | 0.61 | 1.23 | 0.53 |

**Table S7**: Results of the variance analyses of the modeled net assimilation in the SPAC model between the five climates and the three species. The left column corresponds to the outputs of the model with air temperature and acclimated physiology (*i.e.*, based on our measurements on each site). In contrast, the right column presents results obtained with non-acclimated physiological traits (*i.e.*, the traits from the reference site) to isolate the effect of the air temperature. Significant effects ($P < 0.05$) are shown in bold.

| | **ΔT and Acclimation** | | | **ΔT only** | | |
|---|---|---|---|---|---|---|
| | DF | F-value | P-value | DF | F-value | P-value |
| Climate | 4 | 11.261 | **<0.001** | 4 | 4.331 | **0.002** |
| Species | 2 | 23.721 | **<0.001** | 2 | 17.398 | **< 0.001** |
| Climate:Species | 8 | 2.372 | **0.016** | 8 | 0.863 | 0.547 |
| Residuals | 682 | | | 670 | | |

**Table S8**: Results of the Tukey's HSD test of the simulated yearly C budget between the reference and the other sites. The left column corresponds to the outputs of the model with air temperature and acclimated physiology (*i.e.*, based on our measurements on each site), while the right column presents results obtained with non-acclimated physiological traits (*i.e.*, the traits from the reference site) to isolate the effect of the air temperature. Significant effects ($P < 0.05$) are shown in bold.

| | | **ΔT and Acclimation** | **ΔT only** |
|---|---|---|---|
| Species | Relation | *P*-value | *P*-value |
| *T. fortunei* | SubMed - Cold | 0.649224 | 0.999994 |
| | SubMed -Tem | 0.073901 | 0.991281 |
| | SubMed - Med | 0.262958 | 0.975158 |
| | SubMed - SA | **0.04093** | 0.946181 |
| *I. aquifolium* | SubMed - Cold | 0.483926 | 0.402412 |
| | SubMed - Tem | 0.999945 | 0.999971 |
| | SubMed - Med | 0.733006 | 0.999938 |
| | SubMed - SA | 0.11118 | 0.553063 |
| *T. cordata* | SubMed - Cold | **0.040258** | **0.012745** |
| | SubMed - Tem | 0.318164 | 0.891027 |
| | SubMed - Med | **0.023818** | 0.056745 |
| | SubMed - SA | 0.760221 | **0.000103** |

**Table S9**: Results of the Tukey's HSD test of the simulated yearly C budget between the outputs of the model with and without acclimation (*i.e.*, based on our measurements at each site or with the physiologic traits of the reference site, respectively). The variance corresponds to the uncertainty of $J_{max,25}$, and $V_{Cmax,25}$ , $R_{25}$, and $Q_{10}$ measured (n = 37-57). Significant effects ($P < 0.05$) are shown in bold.

| **ΔT & Acclimation *vs*. ΔT only** | | |
|---|---|---|
| Species | Climate | *P*-value |
| *T. fortunei* | Cold | 0.086 |
| | Temperate | **0.015** |
| | Sub-Mediterranean | 0.572 |
| | Mediterranean | **0.046** |
| | Semi-Arid | 0.211 |
| *I. aquifolium* | Cold | 0.480 |
| | Temperate | 0.498 |
| | Sub-Mediterranean | 0.526 |
| | Mediterranean | 0.054 |
| | Semi-Arid | 0.758 |
| *T. cordata* | Cold | 0.163 |
| | Temperate | 0.652 |
| | Sub-Mediterranean | 0.563 |
| | Mediterranean | **<0.001** |
| | Semi-Arid | **0.005** |

### S1. Tree growth, soil humidity, and chlorophyll content

At each campaign, we measured the height of each individual from the basis of the plant to the longest stem. We further measured the leaf chlorophyll content (CC) with a chlorophyll content meter (MC-100; Apogee Instruments; USA). As species-specific leaf anatomical structure can bias the measurements of the MC-100 (Parry, Blonquist and Bugbee, 2014), we calculated correction factors based on the protocol of Weiss (2014). For this, we measured 30 leaves of each species with the MC-100, then collected and stored the same leaves at -80 ºC until lyophilization as in Juillard *et al*. (2024) (Beta 2–8 LD plus; Martin Christ; Germany). We obtain the chlorophyll A and B concentration in µg $ml^{-1}$ through repetitive ethanol dilution and measuring the absorbance of the solution with a spectrophotometer (Synergy Mx; Biotek; USA). We calculated CC per unit leaf area (µmol $m^{-2}$) by multiplying the raw CC by the number of dilutions and dividing by the specific leaf area (SLA) we had measured before lyophilization.

**S2. Functioning of the model**

The model finds an equilibrium for $A_{net}$, $g_S$, and $C_i$ numerically by iterative optimization. Along with physiological and environmental variables, an initial value for $C_i$ and $g_s$ has to be provided for the first iteration. $A_{net}$ is calculated as:

$$A_{net} = g_s\,(C_a - C_i) + R \tag{1}$$

where R is the dark respiration, $C_a$ is the ambient concentration of $CO_2$, and $g_s$ is the stomatal conductance. $g_s$ and $C_i$ are either initial or inherited values. Next, maximum stomatal conductance ($g_{s,\,max}$) is computed with Farquar's equation as:

$$g_{s,max} = \frac{B\,(C_i - \Gamma) - R(vC_i + D)}{(zC_i + D)(C_i - C_a)} \tag{2}$$

where B is the $CO_2$ uptake limiting rate of either electron transport ($J_{max}$) or Rubisco carboxylation ($vc_{max}$), Γ is the $CO_2$ compensation point of photosynthesis, D is a metric of carboxylation and oxygenation rates, and v and z are constants. Next, the model calculates transpiration (E) as:

$$E = gs, max\,\frac{VPD}{P}\,LAI \tag{3}$$

where LAI is the leaf area index. A correction factor ($f_{\psi,leaf}$) for $g_s$ based on leaf water potential ($\psi_{leaf}$) is calculated as:

$$f_{\psi,leaf} = \frac{1 + e^{s_f \psi_f}}{1 + e^{s_f(\psi_f - \psi_{leaf})}} \tag{4}$$

where $\psi_f$ is the reference water potential and $S_f$ is the stomatal sensitivity. Finally, $g_s$ is recalculated following Tuzet's equation:

$$g_s = gmin + \frac{m\,(A_{net} + R)}{C_i - \Gamma} f_{\psi,leaf} \tag{5}$$

where $g_{min}$ is the minimum stomatal conductance, and m is an empirical proportionality factor. Finally, $C_i$ is calculated again from equation 1 and compared to the initial $C_i$. If the difference is important enough, a new iteration is conducted with updated $g_s$ and $C_i$.

### S3. Leaf histology measurements

We measured leaf histological tissue length of 3-10 individuals by producing anatomical cuts as in Didion-Gency *et al.* (2024). First, we clipped two to three leaves of each individual to produce leaf disks 10 mm in diameter. Samples were then fixed in a solution of glutaraldehyde 2.5% and fixed in paraffine with a semi-enclosed benchtop tissue processor (Leica TP1020; Germany) composed of nine tanks of 1L each. The first five tanks were filled with a progressively increasing concentration of alcohol (75, 96, 96, 100, 100%) used to remove water from the samples. The two next tanks were filled with a clearing agent (Ultraclear; Biosystems AG, Switzerland). Finally, samples were put in two tanks filled with melted paraffin. Paraffin-impregnated samples were then fixed in paraffin cubes with an embedding station (HistoCore Arcadia C; Leica; Germany). Finally, we produced slices of 12-20 µm thickness with a microtome (RM 2244; Leica; Germany). After paraffine was removed from the samples with a cleaning agent again, we measured the thickness of histological tissues (lower and upper epidermis, and total leaf thickness (TT)) with a microscope (DMIL; Leica; Germany) equipped with a camera (ICC50HD; Leica; Germany) and the software Leica Application System V4.3 (Leica; Germany) for a total magnification of x400. We calculated the percentage of epidermis (PE) by dividing the sum of epidermises by TT.

Total leaf thickness (TT) varied among sites and species ($P < 0.001$; Figure 8 & Table S2) with leaves of *I. aquifolium* three times thicker than in the other species. Changes in TT among sites differed between species: in *T. fortunei*, leaves were thinner in the cold site than in the sub-Mediterranean, Mediterranean, and semi-arid ones ($P < 0.05$; Table S4). In *I. aquifolium*, leaves tended to get thicker in the semi-arid climate than in the reference sub-Mediterranean site ($P < 0.05$; Table S3). Leaves of *T. cordata* were thinner in the temperate site than in the Mediterranean and semi-arid climates ($P < 0.05$; Table S3).

PE varied between species and sites ($P < 0.001$; Figure 8 & Table S2). *T. cordata* showed the highest PE of the three species, with a mean value of 22.3% while *T. fortunei* and *I. aquifolium* showed respectively 18.3 and 16.3%. In *T. fortunei*, PE increased in cold, temperate, and semi-arid climates compared to the reference site ($P < 0.05$; Table S3).
